# A higher load of deleterious mutations has a detrimental effect on tree growth in maritime pine

**DOI:** 10.1101/2025.11.09.687477

**Authors:** Rosalía Piñeiro, Lowana Cahn, Katharina B. Budde, Agathe Hurel, Adélaïde Theraroz, Giovanni G. Vendramin, Sara Pinosio, Isabelle Lesur, Marina de Miguel, Christophe Plomion, Myriam Heuertz, Santiago C. González-Martínez

**Affiliations:** Evolutionary Biology Research Group (GIBE), Interdisciplinary Centre of Chemistry and Biology (CICA), Universidade da Coruña, Campus da Zapateira sn. 15071 A Coruña, Spain; INRAE, Univ. Bordeaux, BIOGECO, F-33610 Cestas, France; Northwest German Forest Research Institute, Department of Forest Genetic Resources, Professor-Oelkers-Strasse 6, 34346 Hann Münden, Germany; Department of Forest Growth, Silviculture and Genetics, Unit of Genome Research, Austrian Research Centre for Forests, A-1131 Wien, Seckendorff-Gudent-Weg 8, Austria; Institute of Biosciences and BioResources, National Research Council of Italy, Division of Firenze, via Madonna del Piano 10, 50039 Sesto Fiorentino, Italy; EGFV, Univ. Bordeaux, Bordeaux Sciences Agro, INRAE, ISVV, F-33882, Villenave d’Ornon, France

**Keywords:** Purifying selection, genetic load, common garden experiments, forest trees, maladaptive genetic variation, maritime pine, deleterious mutations

## Abstract

A high proportion of deleterious mutations in the genome is predicted to cause a reduction in fitness known as genetic load. In plants, most studies focus on how detrimental mutations increase as a consequence of population bottlenecks, for instance in cultivated species as a negative outcome of domestication or at the margin of species ranges. However, the actual adverse effect of genetic load on individual phenotypes remains little characterised, especially in wild plant species with large populations and long lifespans.

We estimated genetic load based on genomic data in populations of the conifer maritime pine (*Pinus pinaster*, Ait.), and examined its association with phenotypic traits related to growth, phenology, and water-use efficiency measured under common garden conditions.

Our results revealed a strong negative correlation between genetic load and tree height on the Mediterranean island of Corsica, thus suggesting a negative cumulative effect of deleterious mutations on tree growth at the regional scale. Our study is one of the first to experimentally demonstrate adverse phenotypic effects (reduced tree growth) of genetic load in a long-lived plant.

## Introduction

The impact of deleterious mutations on fitness is a relevant topic in evolutionary biology. The cumulative effect of deleterious mutations in the genome is a predicted reduction in individual fitness, modulated by the interaction of several evolutionary forces, including mutation rate, changes in effective population size leading to genetic drift, and selective pressures (Whitlock & Bürger, 2004). Simulation-based studies suggest that deleterious alleles are likely to increase as a consequence of population bottlenecks during range expansions (Gilbert *et al*., 2017; Peischl *et al*., 2018). In contrast, empirical studies validating the impact of deleterious mutations based on experimental evidence are still scarce and constrained mostly to model organisms (Mukai *et al*., 1972; Sandell & Sharp, 2022; Bertorelle *et al*., 2022), particularly for studying recessive genetic diseases in humans (Liu *et al*., 2022). In the case of plants, empirical research efforts are biased towards the study of deleterious mutations in two particular scenarios: cultivated species and recently expanded wild species. The fixation of deleterious variants due to genetic bottlenecks has been observed in a number of crops such as maize, rice or sunflower (Lu *et al*., 2006; Mezmouk & Ross-Ibarra, 2014; Renaut & Rieseberg, 2015; Cornejo *et al*., 2018; Hämälä *et al*., 2021; Conover & Wendel, 2022). This is often associated with the artificial selection of phenotypes of agronomic interest such as fruit size, nutrient contents or resistance to pathogens (Dwivedi *et al*., 2023). However, it might also lead to an overall loss of genetic diversity and reduced fitness (Lye *et al*., 2022; Sun *et al*., 2023) as a negative outcome of the domestication process. In the case of wild plants, research efforts concentrate on the detection of deleterious mutations as a result of demographic changes. Higher proportions of deleterious mutations have been documented in the range margins of recently expanded species. This has been observed in outcrossing herbaceous plants such as *Arabidopsis lyrata*, *Leontodon longirostris* and *Mercurialis annua* (González-Martínez *et al*., 2017; Willi *et al*., 2018; de Pedro *et al*., 2021) but also in a few wild trees (Conte *et al*., 2017), including *Pinus pinea* (the Mediterranean stone pine), thought to have suffered extreme historical bottlenecks (Jaramillo-Correa *et al*., 2020).

Studies that go beyond documenting the increase of deleterious mutations under particular demographic scenarios to show the actual negative impact of genetic load on the plant phenotype are scarce. The adaptive or maladaptive potential of genetic variants can be predicted from genomic sequencing data using computational methods. There are two main approaches: the first is based on sequence annotation (Liu *et al*., 2022), for instance sequence changes introducing stop codons or shifts in the reading frame are highly deleterious. The second is based on the evolutionary conservation of the protein sequence alignments around a mutated site (Choi & Chan, 2015). Mutations in regions that are highly conserved across species are more likely deleterious. In the absence of fitness data, most studies predict the genetic load of plants using computational approaches on genomic data only (Renaut & Rieseberg, 2015; Cornejo *et al*., 2018; Takou *et al*., 2021; Conover & Wendel, 2022). Validation of the actual detrimental effect of these computationally predicted deleterious mutations on the plant phenotype is rare as it requires fitness information from field experiments. This involves taking measurements in common environments, which ensures that the trait variation explored is due to genetic differences by eliminating the effect of the environment, an approach that is particularly time-consuming in long-lived organisms.

In this work, we explored the impact of deleterious mutations on fitness in a long-lived conifer species, maritime pine (*Pinus pinaster* Ait.). Maritime pine shows a natural distribution in the western Mediterranean, Atlantic Europe and North Africa, in warm, temperate oceanic regions. Previous phylogeographic studies, based on biparentally, paternally and maternally inherited neutral markers, revealed a remarkable regional genetic differentiation across its natural range (Burban & Petit, 2003; Naydenov *et al*., 2014; Serra-Varela *et al*., 2015; Jaramillo-Correa *et al*., 2015). This pattern suggests past fragmentation of the species range, probably during the Last Glacial Maximum, followed by limited expansion and relative stability of the divergent gene pools. Our study was performed on a regional scale. We assessed deleterious mutations and fitness in a single-site common garden established in south-western France from seeds collected across the entire natural distribution of maritime pine occurring on the Mediterranean island of Corsica. Corsica provides an ideal setting to study adaptation given the significant ecological diversity associated with the orography of the island, which induces a large span of selective pressures. In addition, on a range-wide scale, provenances representing the entire maritime pine range in Europe and North Africa were investigated in a multi-site clonal common garden experiment planted in Portugal, in North (N), West (W) and Central Spain and in South-West (SW) of France.

Combining genomic and phenotypic data in common environments, this study evaluates the potential effect of deleterious mutations on various fitness components in maritime pine. More specifically, we: (i) estimated the number of deleterious mutations, and thus the genetic load, for individual trees across the distribution range of the species in Corsica, based on either the predicted effect on the protein sequence or the evolutionary conservation of the mutated site; (ii) explored the correlation of deleterious mutations with three phenotypic traits related to fitness: growth, spring phenology and water-use efficiency assessed in common gardens; and (iii) compared the association of genetic load with trait measurements in provenances from Corsica with that of provenances from across the entire species range.

## Materials and Methods

### Sampling of plant material and dataset overview

Our study used plant material representing the maritime pine regional range in Corsica. Phenotypic and genomic (Single Nucleotide Polymorphisms, SNPs) data were generated from maritime pine trees grown from seeds collected in the wild from 30 localities (hereafter provenances; Fig.1; Table 1; Figs. S1 and S2, Table S1a) all across the island in 1994 and 2000 (Durel & Bahrman, 1995) and established in a single-site common garden in Herran, Landes, SW France (hereafter CORSICAPIN common garden). This sampling covered the entire natural distribution of maritime pine in Corsica, spanning a large ecological range from (0-1,200 m a.s.l. and 400-2,000 mm of rainfall/year). CORSICAPIN was planted during three consecutive years in 2010, 2011, and 2012. Each plantation year corresponded to a “plot”. Each plot was in turn made up of five complete blocks, with 20 half-sib families per provenance from the 30 localities randomized within those blocks. Two datasets were generated for the Corsican maritime pine trees: (i) the *CORSICA-capture* dataset, with 133,656 SNPs sequenced on 393 individual trees from 16 Corsican provenances and three phenotypic traits assessed on 177 unrelated individuals (a single individual per family) under common garden conditions; and (ii) the *CORSICA-array* dataset, with 10,185 SNPs genotyped on 334 individuals from 25 Corsican provenances and three phenotypic traits assessed on 334 unrelated individuals (a single individual per family) under common garden conditions. Eleven localities were common for the two datasets of Corsican trees (Fig. 1 and 2; Table 1; Table S1a).

**Figure 1.**
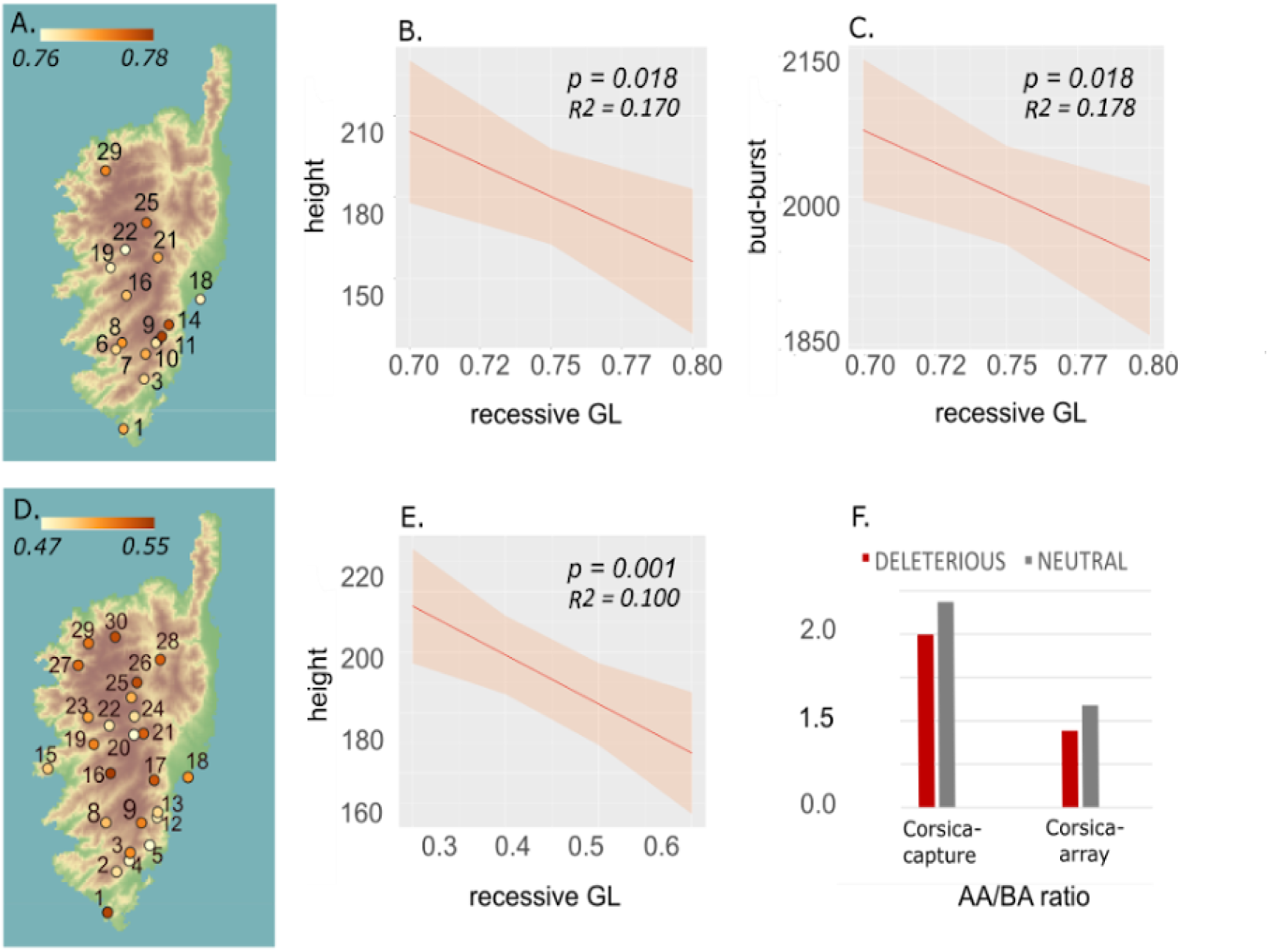
Recessive genetic load of maritime pine across 30 provenances in Corsica. See Table 1 and Table S1a for sampling strategy details. (**A**-**C**) *CORSICA-capture* dataset (393 individuals in 16 provenances): recessive genetic load per locality, A; linear mixed models of phenotypic trait variation with recessive genetic load for tree height in cm, B, and bud-burst time, in accumulated degree-days from the first day of the year, C. (**D-E**) *CORSICA-array* dataset (334 individuals in 25 provenances): recessive genetic load per locality, D; linear mixed models representing the association between recessive genetic load and tree height, E. (**F**) Homozygous-to-heterozygous sites ratios, AA/AB, for deleterious or neutral mutations for the *CORSICA-capture* and *CORSICA-array* datasets. *p*, *p*-value based on conditional F-tests with Kenward-Roger approximation for the degrees of freedom; *R*^2^, conditional goodness of fit (fixed and random effects).

**Figure 2.**
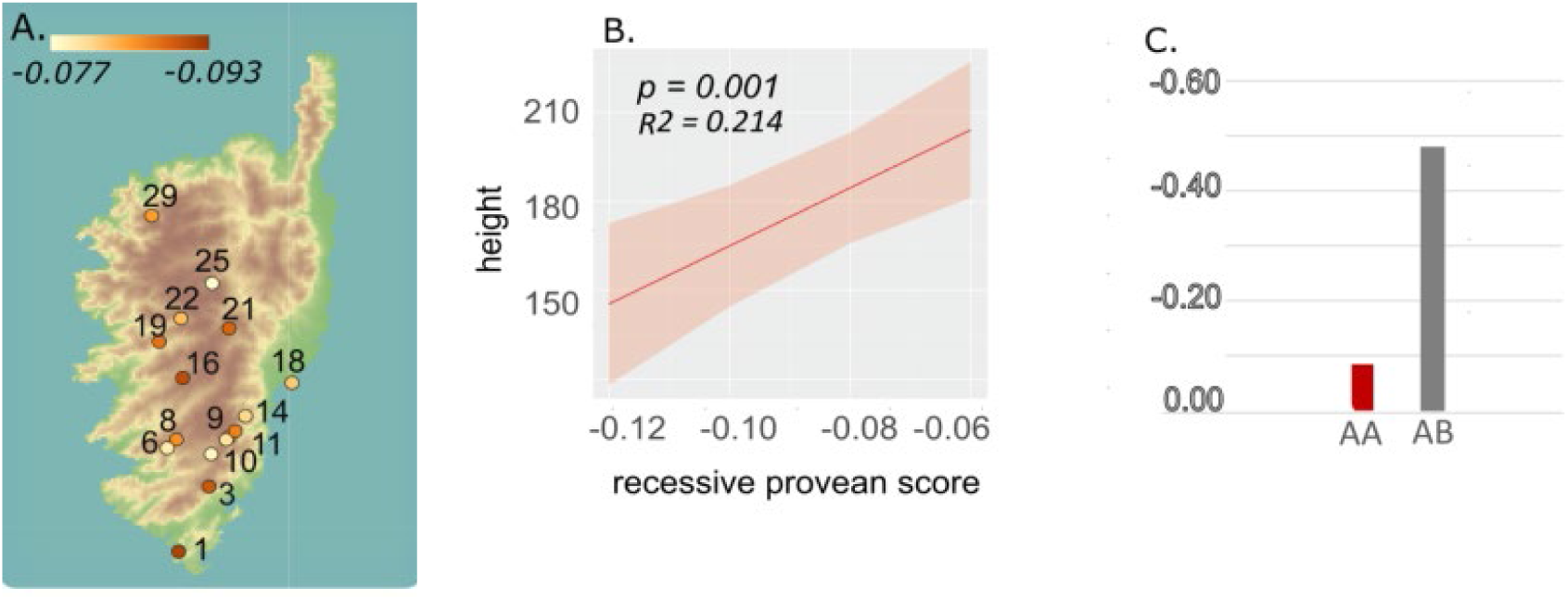
Recessive Provean score (PR) in Corsican populations of maritime pine based on the *CORSICA-capture* dataset (393 individuals from 16 provenances). (**A**) recessive Provean score per locality; (**B**) linear mixed model displaying the correlation between phenotypic trait variation for height (in cm) and recessive PR for tree height; (**C**) average PR at homozygous and heterozygous sites. In B, *p* is the *p*-value based on conditional F-tests with Kenward-Roger approximation for the degrees of freedom; *R*^2^ is the conditional goodness of fit (fixed and random effects).

**Table 1.** Maritime pine phenotypes and DNA datasets included in this study. See Table S1 for details of the sampling strategy and localities.

| *Dataset* | *Phenotypic traits* | *Phenotypic sampling in common garden* | *DNA protocol* | *DNA sampling* | *Geographic scale* |
| --- | --- | --- | --- | --- | --- |
| *CORSICA-capture* | Height, bud-burst, $\delta^{13}\text{C}$ | 177 trees<br>(only unrelated ind.)<br>1-site common garden | Capture<br>133,656 SNPs | 393 ind.<br>16 prov.<br>1 gene pool | Corsica |
| *CORSICA-array* | Height, bud-burst, $\delta^{13}\text{C}$ | 334 trees<br>(only unrelated ind.)<br>1-site common garden | Array<br>10,185 SNPs | 334 ind.<br>25 prov.<br>1 gene pool | Corsica |
| *GLOBAL-array*<br>(de Miguel *et al.*, 2022) | Height, bud-burst, $\delta^{13}\text{C}$,<br>survival | 17,836 trees<br>(464 unrelated ind.;<br>8 clones/ind. per site)<br>5-site clonal garden | Array<br>10,185 SNPs | 464 ind.<br>34 prov.<br>6 gene pools | Southwestern Europe,<br>North Africa |

prov. Provenances; ind. Individuals

We attempted to validate our regional study in Corsica on a range-wide scale by studying maritime pines from 34 additional provenances covering the global European and North African distribution of the species (Fig. 3; Table 1; Fig. S3; Table S1b). These data were obtained from previously published phenotypic and SNP datasets from the CLONAPIN clonal common garden network, a five-site assay containing tree clones from across the entire distribution of the species (de Miguel *et al*., 2022; Archambeau *et al*., 2022). Briefly, collected seeds were germinated in a nursery and one seedling per open-pollinated family was selected to avoid half-siblings (i.e., selected genotypes were unrelated), and vegetatively propagated by cuttings. The maritime pine wide-range dataset (*GLOBAL-array* dataset) consisted of 10,185 SNPs genotyped on 464 unrelated individual trees from 34 provenances and phenotypic traits assessed on 17,836 trees under common garden conditions: eight vegetative clone replicates per provenance and common garden site, and five sites (de Miguel *et al*., 2022; Archambeau *et al*., 2022).

**Figure 3.**
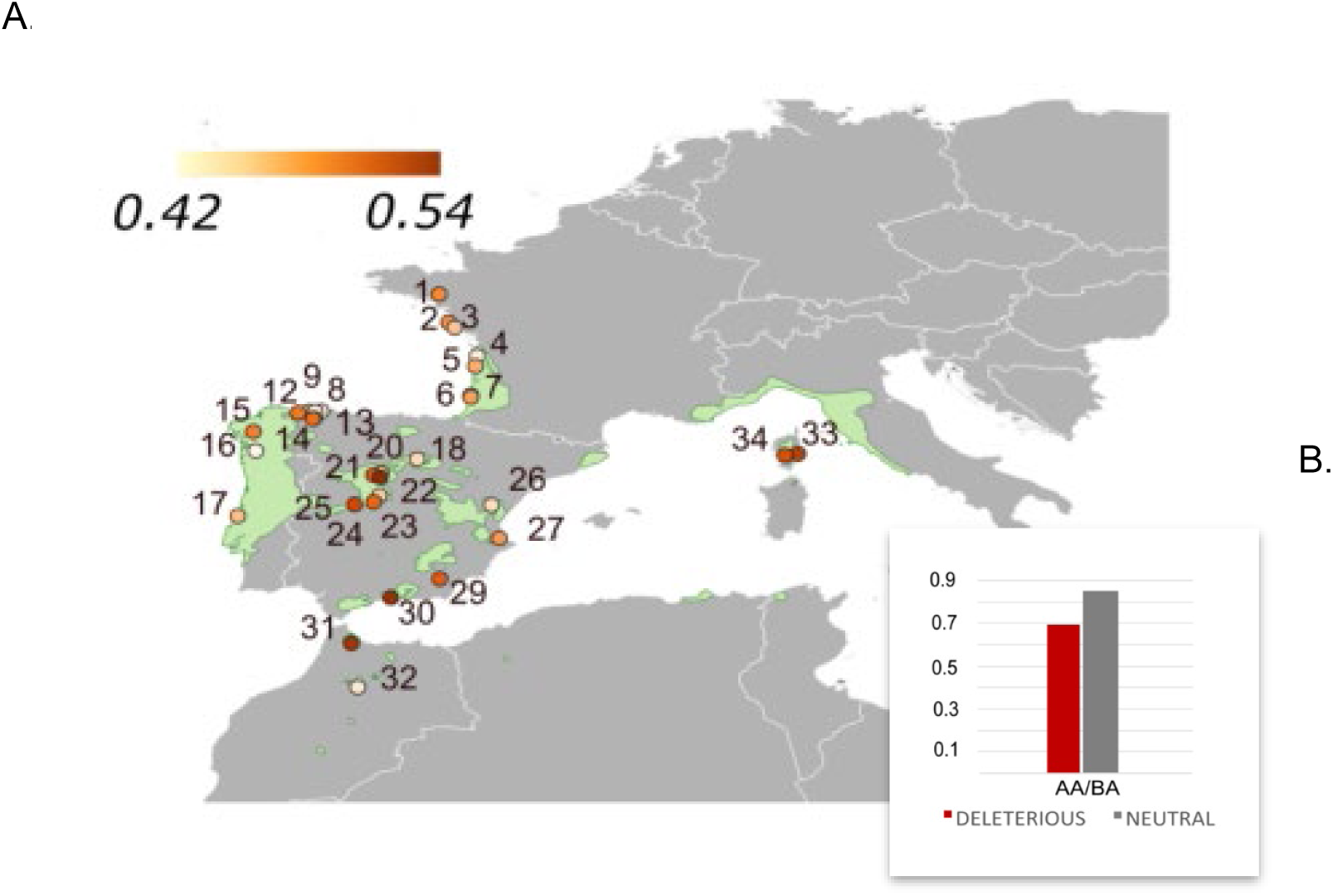
Recessive genetic load across the entire maritime pine range based on the *GLOBAL-array* dataset (N=464 individuals from 34 provenances): (**A**) recessive genetic load per provenance; (**B**) Homozygous-to-heterozygous sites ratio, AA/AB, for deleterious or neutral mutations. See Table 1 and Table S1b for sampling strategy and population details.

### Plant traits used as fitness proxies

To evaluate fitness components related to growth, phenology and drought tolerance, we recorded tree height, spring bud-burst time, and water-use efficiency (WUE) inferred from carbon isotope discrimination (δ^13^C) for each individual tree. Tree height is strongly correlated with growth and early establishment of young forest trees (Bianchi *et al*., 2019). Phenology traits, such as spring bud-burst time, have shown local adaptation along latitudinal or altitudinal clines (Alberto *et al*., 2010), and they might also be associated with resistance to pests and pathogens in forest trees (Krokene *et al*., 2012; Hurel *et al*., 2021). Carbon isotope discrimination can provide integrated measures of water-use efficiency and thus more robust estimates of tree responses to drought than punctual measurements (Plomion *et al*., 2016).

In the CORSICAPIN common garden, tree height was recorded four years after plantation, spring bud-burst time was recorded in 2017 and carbon isotope discrimination (δ^13^C) in 2018. For spring bud-burst, five developmental stages were recorded following (Hurel *et al*., 2021), and stage 3 (S3, brachyblast begins to space) was kept for correlation with genetic load. The Julian day of entry in stage S3 was converted into accumulated degree-days (0°C basis) from the first day of the year, to account for between-year temperature variability. Water-use efficiency (Heschel *et al*., 2002; Donovan *et al*., 2007) was evaluated based on δ^13^C, following standard procedures (Plomion *et al*., 2016). More negative values of δ^13^C indicate lower efficiency. Survival could not be analysed in this common garden as DNA samples were obtained only for surviving trees.

From the five CLONAPIN clonal common gardens, the following raw measurements were retrieved (de Miguel *et al*., 2022): tree height and survival one year after plantation in all five sites, spring bud-burst time in one site (SW France), and carbon isotope discrimination (δ^13^C) in one site (Portugal). These phenotypes represent measurements of 17,836 trees (i.e., eight clones per provenance of each of the 464 unrelated genotypes per common garden site) (Fig. 3; Table 1; Table S1b).

### DNA sequencing and SNP genotyping protocols

New genomic data of Corsican maritime pine trees were obtained using two different protocols, target-capture sequencing yielding 133,656 high-quality SNPs after filtering (*CORSICA-capture* dataset, see below) and SNP-array genotyping of 10,185 SNPs (*CORSICA-array* dataset, including a subset of 2,343 SNPs from the target-capture set). For the *CORSICA-capture* dataset, DNA extraction, target capture and preliminary bioinformatic analysis followed (Milesi *et al*., 2024). Extractions from silica-dried plant material used the DNeasy Plant Mini kit (Qiagen, Hilden, Germany), after a treatment of the samples at -80°C for 24 hours. The capture protocol targeted *Arabidopsis*-orthologous genes and also included candidate genes from previous studies in maritime pine (Jaramillo-Correa *et al*., 2015; Grivet *et al*., 2017). About 3 Mbp of DNA sequence was paired-end sequenced at the IGATech facility (Udine, Italy) on a NovaSeq 6000 (Illumina, San Diego, CA). A Variant Call Format (VCF) was generated in GATK v4.0.10.0 against the loblolly pine (*Pinus taeda* L.) reference genome (v. 2.01, treegenesdb.org). SNPs were filtered to retain only biallelic, highly supported SNPs and remove paralogs, particularly frequent in conifers (Neale *et al*., 2014). First, we discarded low-quality SNPs (QD < 0.25, QUAL < 20, SOR > 3.0, MQ < 30, MQRankSum < -12.5 and ReadPosRankSum < -8.0) and putative paralog-derived SNPs using the HDplot test (heterozygote excess, H, > 0.6, and deviation from the expected read ratio, D, < -20 or D > 20) (McKinney *et al*., 2017). Then, genotype calls of individual trees with sequencing depth (DP) < 8 and genotype quality (GQ) < 20 were recorded as missing. Finally, SNPs and individuals with more than 50% missing calls were filtered out.

For the SNP-array genotyping (*CORSICA-array* and *GLOBAL-array* dataset), we used the multispecies 4TREE Axiom assay (Thermo Fisher Scientific, USA) (Theraroz *et al*., 2024). SNPs were called following the Best Practice Workflow implemented in the Axiom™ Analysis Suite v5.2 and considering dQC ≥ 0.4 and QC call rate ≥ 85%. Following filtering, 10,185 high-quality SNPs were retained. Individuals with more than 30% missing calls were filtered out. The *GLOBAL-array* dataset consisted of the genotypes of 464 tree genotypes from 34 provenances from the CLONAPIN common garden network.

### Population genetic diversity and structure

While the range-wide phylogeography of maritime pine is well studied, the population genetic structure within Corsica has not been investigated with a large set of markers. Previous studies included only a limited number of provenances from the island (e.g., only two provenances in Jaramillo-Correa *et. al.*, 2015) or a limited number of loci (Mariette *et al*., 2001). In our study, we used VCFTOOLS v0.1.16 (Danecek *et al*., 2011) to estimate per-individual observed heterozygosity (*H*_o_), missingness and depth coverage on the *CORSICA-capture* dataset (Fig. S1a). In addition, we performed a Principal Component Analysis (PCA) with SNPrelate (Zheng *et al*., 2012) and a Bayesian clustering with ADMIXTURE v1.23 (Alexander & Lange, 2011) on the same dataset (Fig. S1b,c). The optimal number of genetic clusters (*K*) between 1 and 16 was inferred by the lowest cross-validation error.

### Genetic load estimates from genomic data

We estimated the genetic load for each individual tree based on the *CORSICA-capture* and *CORSICA-array* datasets in two steps. First, we predicted the deleterious effect of each mutation. Second, we summarised the deleterious scores across the genome to estimate the genetic load per individual. Deleterious mutations were predicted using two different approaches: the genetic change at the protein level with SnpEff v. 4.3 (Cingolani *et al*., 2012) and the evolutionary conservation of the protein sequence around mutated sites with Provean v. 1.1.5, which uses ncbi-blast-2.2.29 and cdhit-4.8.1 (Choi & Chan, 2015).

Determination of ancestral and derived allele states was performed for each SNP by comparison with the loblolly pine genome. The effect of each derived mutation on the protein of the exonic regions was predicted with SnpEff, classifying mutations as either neutral or deleterious. The neutral mutations corresponded to derived low-impact mutations not causing amino acid substitutions, while the deleterious mutations included derived deleterious mutations causing amino acid substitutions. When a mutation had several possible effects on the protein, only the deleterious effect was considered for the genetic load calculations. Subsequently, we summarised the predicted deleteriousness scores across the genome into a quantitative genetic load estimate for each individual tree. We used a custom script to compute two genetic load measures per individual tree following (González-Martínez *et al*., 2017): Recessive genetic load (recessive GL; Equation 1a) and additive genetic load (additive GL; Equation 1b), where Nr. Deleterious AA sites is the number of homozygous sites with deleterious mutations, Nr. Neutral AA sites is the number of homozygous sites with neutral mutations, Nr. Deleterious AB sites is the number of heterozygous sites with deleterious mutations, and Nr. Neutral AB sites is the number of heterozygous sites with neutral mutations.

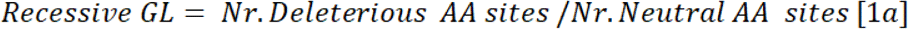

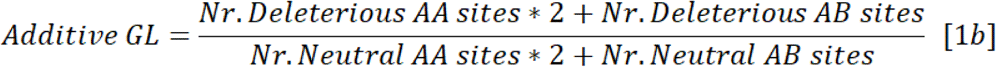

Therefore, the recessive genetic load focuses on the number of homozygous sites containing two deleterious alleles, whereas the additive genetic load indicates the total number of deleterious alleles at homozygous and heterozygous sites. As most deleterious alleles are recessive (Dussex *et al*., 2023), the former ratio is expected to have a higher impact on fitness (González-Martínez *et al*., 2017). Genetic load ratios, i.e., referred to the number of neutral mutations, were used to control for differences in heterozygosity across individuals. It is important to note that per-individual genetic load calculations are not standardised across the plant literature, which reflects the novelty of these approaches.

The effect of mutations was also predicted with Provean, obtaining a quantitative deleteriousness score for each mutation based on evolutionary conservation. First, mutations were classified in two categories, neutral and deleterious, using a threshold for the Provean score at -2.5, following authouŕs recommendations (Choi & Chan, 2015). Thus a score lower than -2.5 corresponds to a deleterious mutation. Second, to avoid using a subjective threshold in the prediction models (Sandell & Sharp, 2022), we averaged the predicted scores across the genome to obtain global scores for each individual tree: an average recessive (recessive PR; Equation 1c) and an average additive (additive PR; Equation 1d) index, where AA sites are the homozygous sites and AB sites are heterozygous sites for derived mutations:

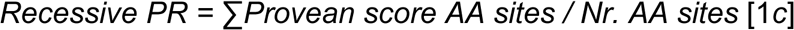

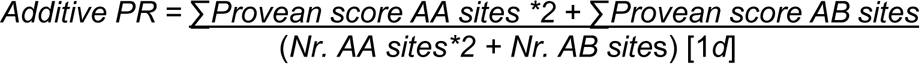

Therefore, the recessive PR indicates the average deleteriousness across derived mutations at homozygous sites whereas the additive PR indicates the average deleteriousness of derived alleles both considering homozygous and heterozygous sites.

### Gene ontology of deleterious mutations

Deleterious mutations identified with SnpEff involved amino-acid changes in 2,233 proteins (see below). With the proteins carrying deleterious mutations identified, we assessed gene ontology (GO) enrichment to find out whether genes carrying deleterious mutations were enriched for specific functions. For all variable sites of the *CORSICA-capture* dataset, corresponding protein sequences were obtained from loblolly pine (v. 2.01, treegenesdb.org) and annotated using PANNZER, Protein ANNotation with Z-scoRE; (Törönen & Holm, 2022) with default parameters. Non-redundant GO terms obtained for proteins containing deleterious mutations were subjected to an enrichment test, comparing them with the full set of GO terms associated with all mutations. For this, we used the topGO R package (Alexa & Rahnenfuhrer, 2024) separately for Biological Processes (BP), Molecular Functions (MF) and Cellular Components (CC) root categories and Fisher exact tests. Finally, GO subgraphs of node size 10 (i.e., including the 10 most significant GO terms in the Fisher tests) were produced for each root category.

### Statistical correlation of genetic load with fitness proxies

We investigated whether the accumulation of the deleterious mutations per individual tree predicted above had an actual detrimental effect on the phenotype. To this end, we explored the correlation between phenotypic data obtained in common gardens, used as fitness proxies, and the genetic load (GL) or Provean scores (PR) estimated from genomic data with SnpEff and Provean, respectively. We fitted linear mixed-effects models in R using the lmer/glmer functions in package lme4 (Bates *et al*., 2015). For the Corsican datasets (*CORSICA-capture* and *CORSICA-array*), we used the plot and block within plot as random effects (Equation 2a), as the trees in the different plots were planted in different years (see above) and no clones were used.

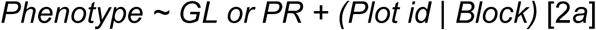

In this way, we controlled for age differences across plots for the traits measured in the same year (bud-burst time at stage 3, i.e., S3, and δ^13^C) and for differences across years for total height, measured at the same age but in different years, as well as for spatial microenvironmental variation in both cases (i.e., the block effect). In cases where the random effect of block exhibited zero variance (height and bud-burst in the *CORSICA-capture* dataset for both additive and recessive genetic load, and δ^13^C with additive genetic load in the *CORSICA-array* data), we dropped it from the model to avoid overfitting. Similarly, we did not include any random effect for provenance considering the lack of genetic structure in the dataset (see below). To maintain the independence of genotypes, we kept the estimates of genetic load only for one individual per family (see above). Either we selected the individual with the highest genotype quality (i.e., with the fewest missing data), or we selected it randomly when quality was similar. As tree height has been found to decrease with altitude in Corsica (Durel & Bahrman, 1995; Hurel *et al*., 2021), the mean altitude per original provenance was also accounted for as a random effect in a second series of models (Equation 2b).

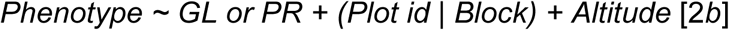

For the range-wide dataset (*GLOBAL-array* dataset), we first used the multisite phenotypic measurements of tree height and survival. We considered the block within site (to control for differences between different common garden sites and the spatial microenvironmental variation within sites), and the clones nested in populations (to control for the genetic differences among clones within provenances of origin) as random effects (Equation 3a). A previous study on the five CLONAPIN common gardens, showed that the genotype-per-environment interaction, GxE, explained only 1.5% of the variance in height, being also not significantly different from zero (see model M2 and Table S5 in Archambeau *et al*., 2022). Thus, we considered the GxE interaction to be negligible for our analyses. Single common garden sites in SW France and Portugal were used to measure spring bud-burst time and δ^13^C, respectively (Equation 4a). The population genetic structure across the species’ range was accounted for by using regional gene pools (Atlantic France, Atlantic Iberia, Central Spain, Southeastern Spain, Corsica, and Morocco; Fig. S3) as random effects (Jaramillo-Correa *et al*., 2015). This analysis was repeated excluding admixed individuals (i.e., those with STRUCTURE assignments to a single gene pool <0.70).

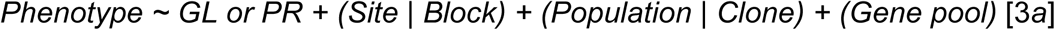

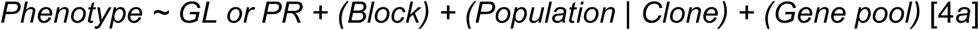

Models including altitude nested to gene pool as a random effect were also explored (Equations 3b and 4b).

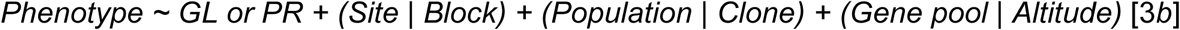

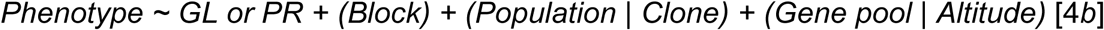

For continuous phenotypic traits (see above), we verified that their distributions followed a normal distribution and then fitted linear models using the lmer function. Survival was treated as a categorical variable and then fitted generalised linear mixed-effects models using the glmer function. For each of the four phenotypic traits studied (height, spring bud-burst time, δ^13^C and survival), correlations were calculated with both the recessive and additive genetic load, as defined above. Control parameters were defined using lmerControl in the R package lme4, keeping default parameters except for the gradient of the deviance function for convergence that was set to a tolerance of 0.0025.

## Results

### DNA sequencing and SNP genotyping

After filtering low-quality SNPs, paralogs and missing data, the final *CORSICA-capture* and *CORSICA-array* datasets contained 133,656 and 10,185 SNPs genotyped in 177 and 334 individuals, respectively. The latter including a subset of 2.343 SNPs from the former.

### Population genetic diversity and structure

The extensive DNA sequencing used in the *CORSICA-capture* dataset revealed lack of genetic divergence between 16 maritime pine localities in Corsica, which has a total surface of 8,681 km^2^, ensuring that there is no hidden genetic structure that might blur the signals of natural selection, an important source of bias crucial to consider when studying local adaptation (Beaumont & Balding, 2004; Williamson *et al*., 2005; Barton *et al*., 2019). No differences in levels of observed heterozygosity were revealed in individuals across island localities, despite the ecological differentiation associated with the complex orography of Corsica, where a central mountain chain runs its length and three forested ecoregions are distinguished (Institut National de l’information géographique et forestière, IGN, https://inventaire-forestier.ign.fr/). Per-individual observed heterozygosity (*H*_o_) ranged from 0.03 to 0.07 (average 0.05; median 0.05) and did not display any evident geographic pattern (Fig. S1a). Missingness per individual was, on average 0.22 (range 0.03-0.52; median 0.19) and individual depth coverage was 21x (range 11x-45x; median 20x). In addition, little genetic structure was revealed within the island, both by the PCA and by the high proportion of individuals with admixed ancestry estimated by the Bayesian clustering analyses. The results of the admixture analysis in Corsica showed an optimum at *K*=1, where the cross-validation error was minimised, revealing a weak geographic structure of the genetic variation (Fig. S1b). At *K*=2 to *K*=16, the groups were not in agreement with the geography and many individuals exhibited ancestry from two or three genetic groups. PC1 explained just 1.14% of the total genetic variance, while PC2 and PC3 explained 0.83% and 0.55%, respectively (Figure S1c). The scatterplot on the three principal components showed that most individuals widely overlaid, while a slight differentiation of individuals from Ventilègne (a relatively isolated population at the southernmost part of the island) was observed, in agreement with the low levels of admixture detected in this locality by the genetic clustering analysis.

### Estimates of genetic load and Provean scores from genomic datasets

Based on SnpEff, the mutation impact was annotated for the *CORSICA-capture* dataset in 44,688 exonic SNPs: 28,298 as deleterious involving amino acid changes in 2,233 proteins, and 16,390 as neutral. For the *CORSICA-array* dataset, 1,324 SNPs were annotated: 478 as deleterious and 846 as neutral. When mutations had several impacts, the deleterious one was considered, involving only 0.06% of the annotated SNPs in both datasets. Per-individual recessive GL scaled by the number of neutral mutations was, on average, 0.74 for the capture and 0.44 for the array dataset, while the additive GL was 0.77 for the capture and 0.49 for the array dataset. No differences across individuals from different localities were observed (Fig. 1a,d; Fig. S2). Per-individual counts of sites with deleterious mutations, revealed different trends in homozygosity and heterozygosity (Fig. 1f). For the *CORSICA-capture*, the average homozygous-to-heterozygous sites ratio per individual was 1.99 for deleterious mutations and 2.37 for neutral mutations. For the *CORSICA-array* dataset, it was also smaller for deleterious (0.89) than for neutral (1.18) mutations.

Based on Provean, 39,182 SNPs were annotated for the *CORSICA-capture* dataset: 8,523 deleterious and 30,659 neutral. For the *CORSICA-array* dataset, 47 SNPs were annotated as deleterious and 932 as neutral. The average per-individual Provean scores did not show any differences across localities (Fig. 2a; Fig. S2). Provean scores also revealed different trends in homozygosity and heterozygosity (Fig. 2c), showing a larger average, i.e., less deleterious, average Provean scores at homozygous sites compared to heterozygous sites both for *CORSICA-capture* (-0.08 vs -0.48) and *CORSICA-array* (-0.17 vs -0.24) datasets. This can be interpreted as natural selection preferentially removing more deleterious mutations (with more negative Provean scores) at homozygous sites.

The results of the GO enrichment (Fig. S4) showed that genes carrying deleterious mutations were enriched for 14 Biological Processes (BP), 15 Molecular Functions (MF) and six Cellular Components (CC) essential to plant and cell functions. Importantly, genes involved in the BP process “biosynthesis of fatty acids” and the MF “symporter activity” showed highly-significant enrichment of deleterious mutations predicted both by Provean and SnpEff. Fatty acid biosynthesis in plants is a crucial metabolic process that results in the formation of membrane lipids and storage oils. Symporter activity plays a crucial role in plant nutrient uptake, transport, and distribution (Vaughn et al., 2002; Zhang et al., 2024).

### Correlation of genetic load and Provean scores with fitness proxies on a regional scale

The analyses exploring the association of phenotypic and genetic load variation across Corsican individuals revealed a negative correlation of the SnpEff-estimated genetic load with the phenotypic traits related to tree growth in all the analyses and with spring bud-burst time in some of the analyses. For the *CORSICA-capture* dataset (Fig. 1b,c; Table S2a), the recessive GL showed a significant correlation with tree height and with spring bud-burst variation, while the additive GL was only marginally correlated with tree height and significantly correlated with spring bud-burst variation. For the *CORSICA-array* dataset (Fig. 1e; Table S2b), both the recessive GL and the additive GL showed a strong and significant correlation with tree height while no correlation with spring bud-burst was detected. Water use efficiency as, estimated by δ^13^C, a trait that may be adaptive in drought conditions (Plomion *et al*., 2016), only showed a marginally significant effect with the recessive and additive GL for the *CORSICA-capture* dataset (Table S2a), whereas it did not exhibit any significant correlations for the *CORSICA-array* dataset (Table S2b).

When the altitude of provenance was accounted for as a random effect, we observed a significant contribution of this factor to the variation of tree height in Corsica. However, when controlling for altitude effect in the *CORSICA-capture* dataset, the negative correlations of recessive GL with tree height and spring bud-burst and of additive GL with spring bud-burst remained significant (Table S2a). In the *CORSICA-array* dataset, negative correlations of the recessive and additive genetic load with tree height also remained significant (Table S2b). Altitude showed zero variance on water use efficiency and was thus not included in the analysis (results not shown).

Provean predictions confirmed a significant reduction of tree height with increasing deleteriousness in recessive PR (more negative recessive PR) for the *CORSICA-capture* dataset (Fig. 2b; Table S3a) and a marginally significant decline for the *CORSICA-array* dataset (Table S3b), while no significant height reduction was found with the additive PR. When the altitude of provenance was accounted for as a random effect, correlations remained virtually unchanged. Water use efficiency δ^13^C and spring bud-burst did not exhibit any significant correlations (Tables S3a,b) with either recessive or additive PR.

### Validation of genetic load correlation with fitness proxies on a range-wide scale

In the case of the *GLOBAL-array* dataset, 478 SNPs were annotated as deleterious with SnpEff and only 47 SNPs with Provean. For SnpEff predictions, the *GLOBAL-array* dataset exhibited an average recessive GL of 0.47 and additive load of 0.51 per individual and no clear geographic trends (Fig. 3a; Fig. S3, as in the Corsican datasets. The average homozygous-to-heterozygous sites ratio per individual was also lower for deleterious mutations than for neutral mutations (0.69<0.85; Fig. 3b). Provean scores in the *GLOBAL-array* also revealed larger, i.e., less deleterious, average Provean scores at homozygous sites compared to heterozygous sites (-0.14 vs -0.23), as for the analyses at the regional scale in Corsica.

The analyses exploring the phenotypic and genetic load variation across range-wide individuals revealed a negative correlation of the genetic load with tree growth, as in the Corsican datasets. The recessive GL estimated using SnpEff was negatively correlated with tree height (Table 2; Table S2c), while the additive GL was only marginally correlated with tree height (Table S2c). No correlations between recessive or additive GL, and spring bud-burst, δ^13^C or survival were detected (Table S2c). When using Provean, the association of tree height with the recessive PR was not significant (Table S3c), while tree height was found to significantly increase with lower (more deleterious) additive PR. When altitude of the provenance was accounted for, the associations remained virtually unchanged (Table S3c). These results are probably due to the low number of annotations obtained with Provean for the *GLOBAL-array* dataset.

**Table 2.** Linear mixed models of the association of tree height with recessive genetic load (GL) in the range-wide maritime pine *GLOBAL-array* dataset.

| *Predictors* | *Estimates* | *CI* | *p* |
| --- | --- | --- | --- |
| (Intercept) | 378.91 | 170.58 – 587.24 | **<0.001** |
| recessive GL | -94.31 | -178.85 – -9.77 | **0.029** |
| Observations | 11029 |  |  |
| Marg / Cond $R^2$ | 0.000 / 0.774 | | |

CI, confidence intervals; *p*, *p*-value based on conditional F-tests with Kenward-Roger approximation for the degrees of freedom; Marg $R^2$ marginal goodness of fit (fixed effects); Cond $R^2$ , conditional goodness of fit (fixed and random effects). See Table S2c for details on random effects.

## Discussion

This work investigates the effect of deleterious mutations on maritime pine fitness by combining genomic data and fitness proxies estimated from phenotypic traits measured under common garden conditions. Our results revealed a genetic signal of ongoing purifying selection and a significant decline of total tree height with genetic load, which suggests a detrimental cumulative effect of deleterious mutations.

### Genomic signatures of purifying selection in Corsica

Computational methods used for predicting deleterious mutations from genomic data revealed a genetic signature of purifying selection in maritime pine in two different datasets of Corsican trees. Genetic load estimates revealed different trends in homozygosity and heterozygosity, suggesting that natural selection is preferentially removing deleterious mutations at homozygous sites. This is indicated by the lower homozygous-to-heterozygous sites ratios found at sites with mutations with a deleterious effect on the protein predicted with SnpEff compared to sites with neutral mutations. Provean, based on the evolutionary conservation of the gene regions bearing mutated sites, confirmed this result by predicting larger **—**i.e., less deleterious**—** scores at homozygous sites compared to heterozygous sites.

Purifying selection is the most prevalent form of selection (Agrawal & Whitlock, 2012). Under purifying selection deleterious mutations at homozygous sites generate individual phenotypes with reduced fitness that are purged in each generation, while deleterious mutations at heterozygous sites, at least at loci with recessive deleterious alleles, may escape purifying selection and accumulate in a manner more similar to neutral mutations. The preferential removal of recessive mutations at homozygous sites has been described by theoretical models (Peischl *et al*., 2018) and is also widely observed empirically in genomic studies. This includes the model animal species *Drosophila melanogaster* (Mukai *et al*., 1972) as well as non-model species, mostly vertebrates, such as killer whales, snub-nosed monkeys, great apes or sea otters (de Valles-Ibáñez *et al*., 2016; Beichman *et al*., 2019; Kuang *et al*., 2020; Foote *et al*., 2021) but also a number of model and non-model plants such as maize (Mezmouk & Ross-Ibarra, 2014), the model tree *Populus trichocarpa* (Zhang *et al*., 2016), sunflower, cotton, *Arabidopsis lyrata* or *Mercurialis annua* (Renaut & Rieseberg, 2015; González-Martínez *et al*., 2017; Takou *et al*., 2021; Conover & Wendel, 2022).

Under scenarios of demographic changes, such as population expansions and contractions, genetic drift becomes the predominant evolutionary force leading to a relaxation of purifying selection. This might lead to the increase of genetic load at the edges of the species’ distribution range, caused by the inefficient removal of deleterious variants. In the case of the Corsican maritime pine, we observed no differences between the genetic load or Provean scores in peripheral and central localities, which suggests no relaxation of purifying selection occurred at the island margins. The absence of divergent lineages of maritime pine within Corsica revealed by Bayesian clustering along with the similar levels of genetic diversity and genetic load found in individuals from range-margin and central localities of the island, supports that maritime pine might be functioning as a panmictic population in Corsica, discarding a scenario of recent expansion and relaxed purifying selection. In agreement with this observation, in the *GLOBAL-array* dataset, the genetic load in Corsica showed genetic load values comparable to the continental localities. However, this dataset does not include populations along the Mediterranean coast of France and northern Italy, known to be the closest to the Corsican ones based on genetic data (Naydenov *et al*., 2014). The Corsican palynological record indicates that maritime pine was already present on the island ca. 2,000 years ago (Carcaillet *et al*., 1997).

### Negative impact of deleterious variants on maritime pine fitness in Corsica

By combining genomic data and fitness proxies, our results revealed a decline of tree height with genetic load in Corsica. This decline was significant for the two datasets with Corsican trees based on SnpEff predictions of the recessive genetic load and, for the capture dataset (marginally significant for the array dataset), and also based on Provean recessive scores. This points to a detrimental cumulative effect of the deleterious mutations in homozygosis on tree growth. Tree height exhibited a large variation within and between provenances in Corsica, without any clear geographical trend, which suggests that our common garden experiment was suitable for fully exploring all the genetically determined variability on this trait. This is probably due to the heterogeneous conditions in the local environment in terms of altitude, rainfall or temperature. In contrast to the recessive genetic load, which focuses on deleterious homozygous sites, significant associations between tree height and the additive genetic load were only detected for one of the Corsican datasets based on SnpEff while no significant associations were detected in any of the two datasets based on Provean. This suggests a predominance of the effect of the “realised” genetic load at homozygous sites (Bertorelle *et al*., 2022).

Population genetics studies on model and non-model organisms have substantially contributed to our understanding of how genetic load manifests in genomes. However, studies connecting genome-wide bioinformatic predictions and experimental validation of their effect on fitness are still rare in plants. To our knowledge, effects on traits of predicted genetic load have been documented previously in a study on the North American red spruce (Capblancq *et al*., 2021), where negative impacts of genetic load on tree growth were also detected. However, that study involved only a 12-week common garden experiment. Other studies testing the effect of genetic load on fitness were either performed under non-common garden conditions (Zhang *et al*., 2016), did not find any significant effects, e.g. *Arabidopsis lyrata* (Takou *et al*., 2021), or were developed in the context of hybridisation (Conte *et al*., 2017). To date, approaches combining genomic assessments and fitness measurements in experimental conditions have prioritised the identification of favourable genetic variants under positive selection in natural tree populations (Savolainen *et al*., 2007) rather than detrimental mutations.

A negative association of spring phenology (bud-burst) with genetic load was only detected for one of the Corsican datasets based on SnpEff predictions, which might indicate a less straightforward relationship between phenology and tree fitness. The importance of bud-burst for fitness might be related to the avoidance of unfavourable environmental conditions while maximising exposure to favourable conditions during the growing season (Bennie *et al*., 2010). Too early emergence might expose leaves to damaging frost or foliar pests while late emergence might prevent the full utilisation of the growing season. Bud-burst time has shown local adaptation along latitudinal or altitudinal clines and might also show asynchrony with the current changing climate (Alberto *et al*., 2010), and both tree height and spring bud-burst time also seem to be associated with resistance to pests and pathogens (Sampaio *et al*., 2016; Hurel *et al*., 2021).

The absence of a correlation of the genetic load with water-use efficiency, might be due to a limitation of our common garden experimental (Hurel, 2019), as the high annual mean precipitation (∼1000 mm) and mild temperatures of the CORSICAPIN common garden limit the exploration of water-use efficiency using δ^13^C. Water-use efficient individuals are expected to acquire more biomass per unit of water transpired and typically enjoy greater biomass accumulation and fitness (Heschel *et al*., 2002; Donovan *et al*., 2007), but only under drought or arid conditions.

### Validation on a range-wide geographical scale

The regional-scale genetic load estimates were validated on a range-wide scale by re-analysing genetic data from 34 provenances across Europe and North Africa in the *GLOBAL-array* dataset. These analyses partially confirmed the Corsican ones. First, computational predictions showed a lower homozygous-to-heterozygous sites ratio at sites with deleterious mutations relative to sites with neutral mutations predicted with SnpEff and larger (less deleterious) average Provean scores at homozygous sites compared to heterozygous sites, which suggests purifying selection acts on recessive deleterious mutations. Second, per-individual genetic load and Provean scores did not significantly differ between different regions or central-range populations. Finally, the significant decline of tree height with recessive genetic load was found using SnpEff predictions but not with Provean scores. This result is likely reflecting the lower number of deleterious sites predicted by Provean compared to SnpEff, which adds up to the relatively low sampling sizes per gene pool in the *GLOBAL-array* dataset. In large panmictic populations or metapopulations connected by gene flow, such as it is typical in maritime pine, deleterious alleles are expected to be held at extremely low frequencies by purifying selection, so that they can remain undetected under low sampling efforts.

Altogether, our results point to a significant detrimental cumulative effect of deleterious mutations on tree growth in maritime pine. This finding provides innovative insights into the genetic basis of adaptation of forest trees to their environment, with the aim of incorporating the control of deleterious mutations into breeding and conservation programmes, which is particularly relevant in the current context of acute population vulnerability of forest tree species imposed by climate change.

## Supporting information

Figures S1 to S3 Tables S1 to S2 SI References

## Acknowledgements

We thank S. Scalabrin, J. Archambeau, D. Macaya-Sanz and L. Duvaux for genetic data production and sampling. We also thank A. Saldaña, F. del Caño, E. Ballesteros and D. Barba (ICIFOR, INIA, Madrid) and the ‘Unité Expérimentale Forêt Pierroton’ (UEFP, INRAE, Cestas; doi: 10.15454/1.5483264699193726E12) for common garden establishment, measurements and sampling help, as well as the Genotoul bioinformatics platform Toulouse Midi-Pyrenees and the Centro de Supercomputación de Galicia (CESGA) for computing resources. This study was funded by the IdEX Bordeaux University ‘Investments for the Future’ program / GPR Bordeaux Plant Sciences, the ‘Chaires d’installation 2015’ (EcoGenPin), the INRAE ECODIV Division (project POSTD2020-8), and the María Zambrano talent-attraction programme (Spanish Ministry of Innovation and Universities).

## Competing interests

The authors do not have organizational affiliations, stock, patent filings or research support to disclose

## Author Contributions

S.C.G.M., and R.P. designed the research.

C.P. established the experiments (common gardens).

S.P., G.G.V., A.H., A.T., and M.D.M. generated and curated the data

R.P., L.C., S.C.G.M, K.B.B., and I.L. analysed the data.

R.P., S.C.G.M., K.B.B., and M.H. wrote the paper.

S.C.G.M, M.H., and C.P. coordinated the associated projects.

## Data availability

Two genomic datasets for the Corsican maritime pine trees are provided: 1) the *CORSICA-capture* dataset (vcf format), with 133,656 SNPs and 393 individual trees from 16 localities across the island (including Provean and SnpEff annotations); 2) the *CORSICA-array* dataset (popgen format), with 10,185 SNPs and 334 individuals from 25 different localities (https://doi.org/10.57745/N9F7DA).

## Notes

https://doi.org/10.57745/N9F7DA)

### Competing Interest Statement

The authors have declared no competing interest.

