## Supplementary material for "A higher load of deleterious mutations has a detrimental effect on tree growth in maritime pine": Figures S1 to S3 Tables S1 to S2 SI References

* *Rosalía Piñeiro*

**This PDF file includes:**

Figures S1 to S3
Tables S1 to S2
SI References

**Fig. S1.** Genetic diversity and population structure of maritime pine in Corsica, inferred from the *CORSICA-capture* dataset (A) Average per-individual genetic diversity per provenance (B) The results of the Admixture analysis in Corsica showed that from K=2 to K= 16 the groups do not fit the geography and most individuals exhibited ancestry from two or three genetic groups. Also, we found an optimum number of gene pools (K) at K=1, where a minimum cross-validation value was reached, revealing an apparent lack of geographic structure of the genetic variation. (C) The PCA also revealed a weak genetic structure, with PC1 explaining 1.14% of the genetic variance, and PC2 and PC3 0.83% and 0.55%, respectively. The scatterplots on the three principal components show that most individuals widely overlay. See Table S1 for locality details.

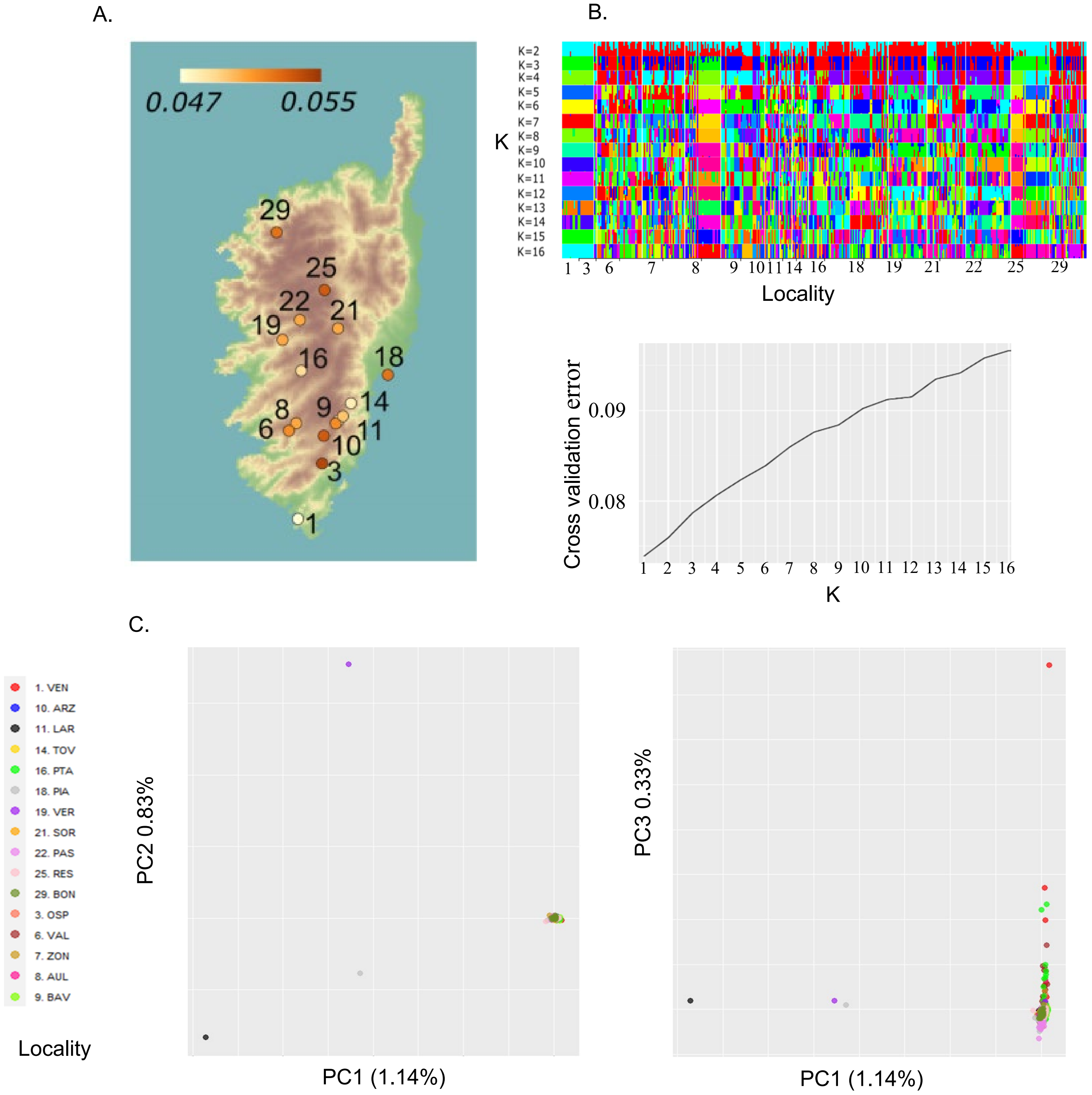

**Fig. S2.** Average per-individual genetic load in each provenance of maritime pine in Corsica. *CORSICA-capture* dataset (N=393 individuals from 16 provenances): (A) recessive genetic load, (B) additive genetic load. *CORSICA-array* dataset (N=334 individuals from 25 provenances): (C) recessive genetic load, (D) additive genetic load. See Table S1 for locality details.

Genetic Load | SnpEff

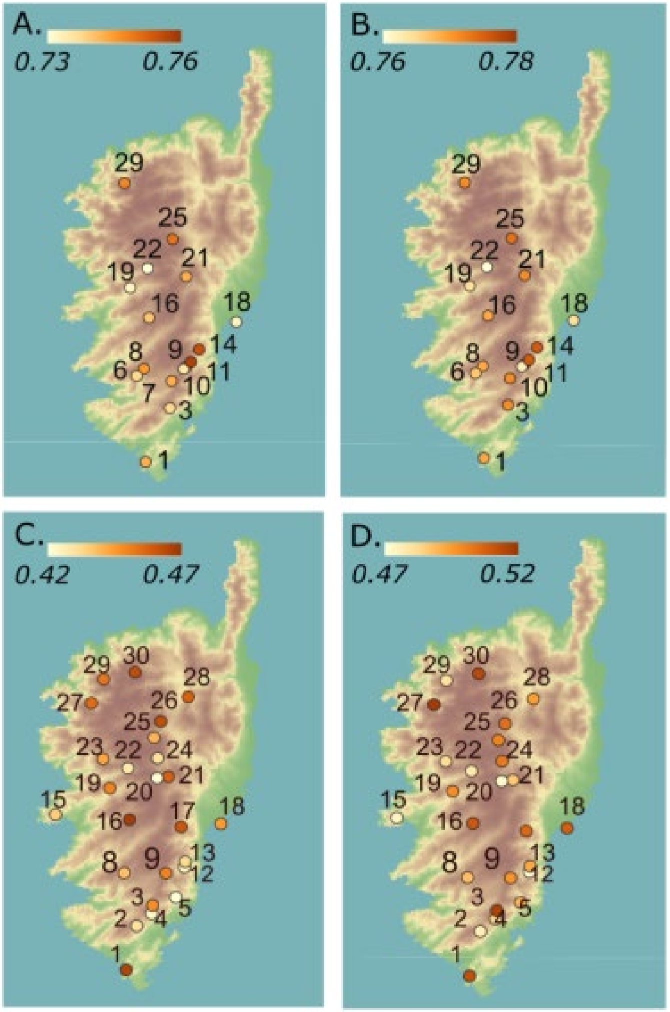

**Fig. S2 (cont.).** Average Provean scores per-individual in each provenance of maritime pine in Corsica. *CORSICA-capture* dataset (N=393 individuals from 16 provenances): (A) recessive average Provean score, (B) additive average Provean score. *CORSICA-array* dataset (N=334 individuals from 25 provenances): (C) recessive average Provean score, (D) additive average Provean score. See Table S1 for locality details.

Average scores | Provean

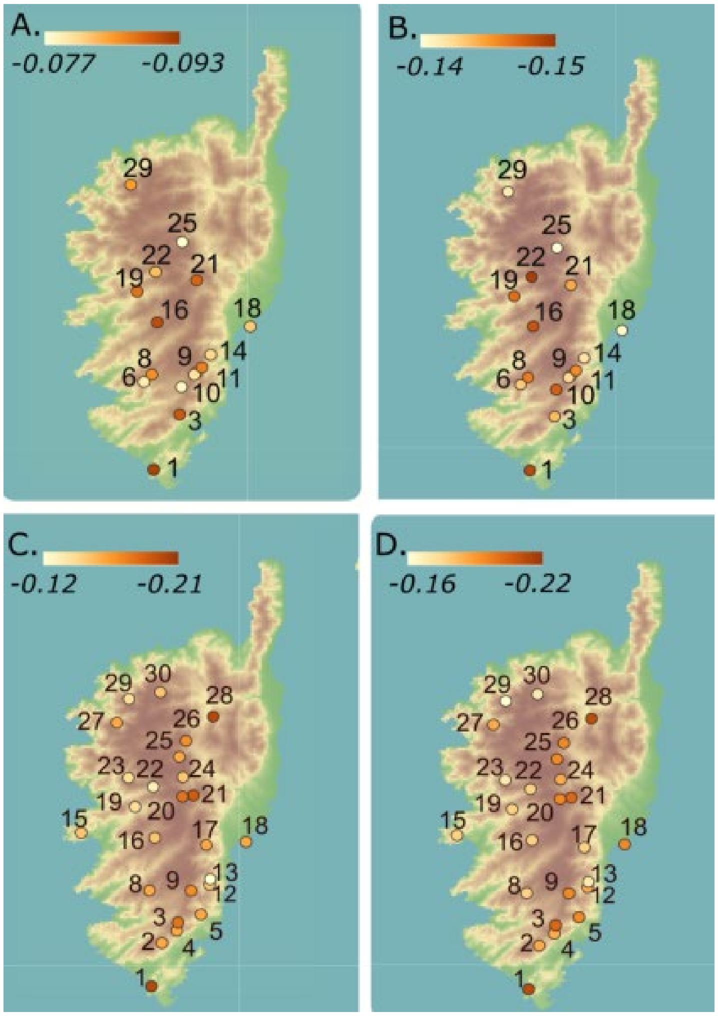

**Fig. S3.** Average per-individual genetic load in each locality across the entire maritime pine range in Europe and North Africa based on the *GLOBAL-array* dataset (N=464 individuals from 34 provenances): (A) recessive genetic load, (B) additive genetic load, (C) recessive average Provean score, (D) additive average Provean score. The distribution range of maritime pine is outlined in green. (E) Distribution of the six divergent maritime pine gene pools -outlined in different colours- modified from de-Miguel *et al*. (2022). See Table S1 for population details.

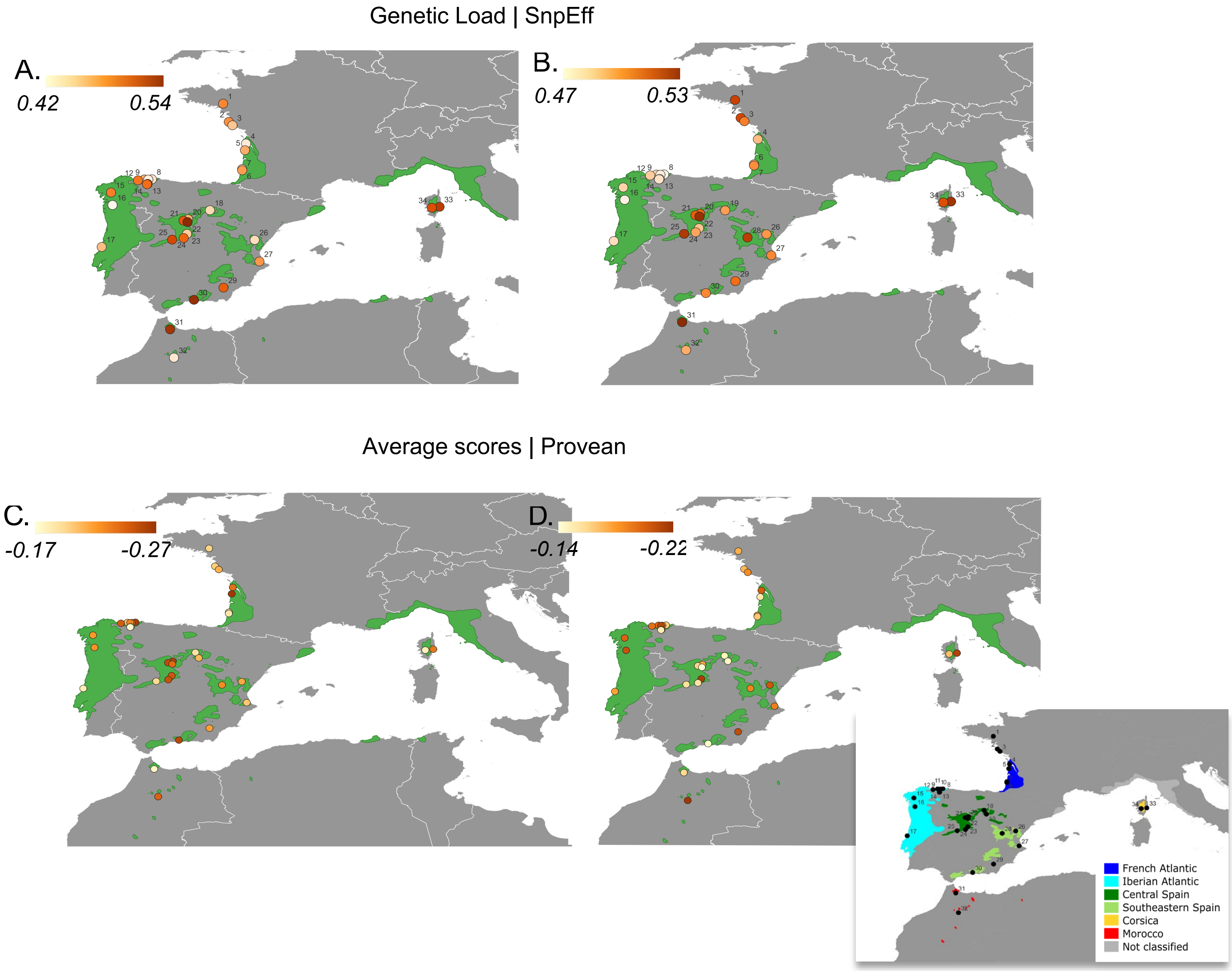

**Fig. S4.** Gene ontology enrichment for genes carrying deleterious mutations enriched for specific functions in maritime pine for the CORSICA-*capture* dataset based on SnpEff and Provean. Three root categories were considered: Biological Processes - BP, Cellular Components- CC, and Molecular Functions - MF. For each category, (A) the ten most significant GO terms in the Fisher tests for each root category and (B) the GO subgraphs of node size 10 (i.e., including the 10 most significant GO terms in the Fisher tests) were produced. Only high-quality annotations were considered (ARGOT_PPV > 0.5). Significant enrichment is indicated in red (p<0.01) or purple (p<0.05).

A.

Biological Processes - BP | SnpEff

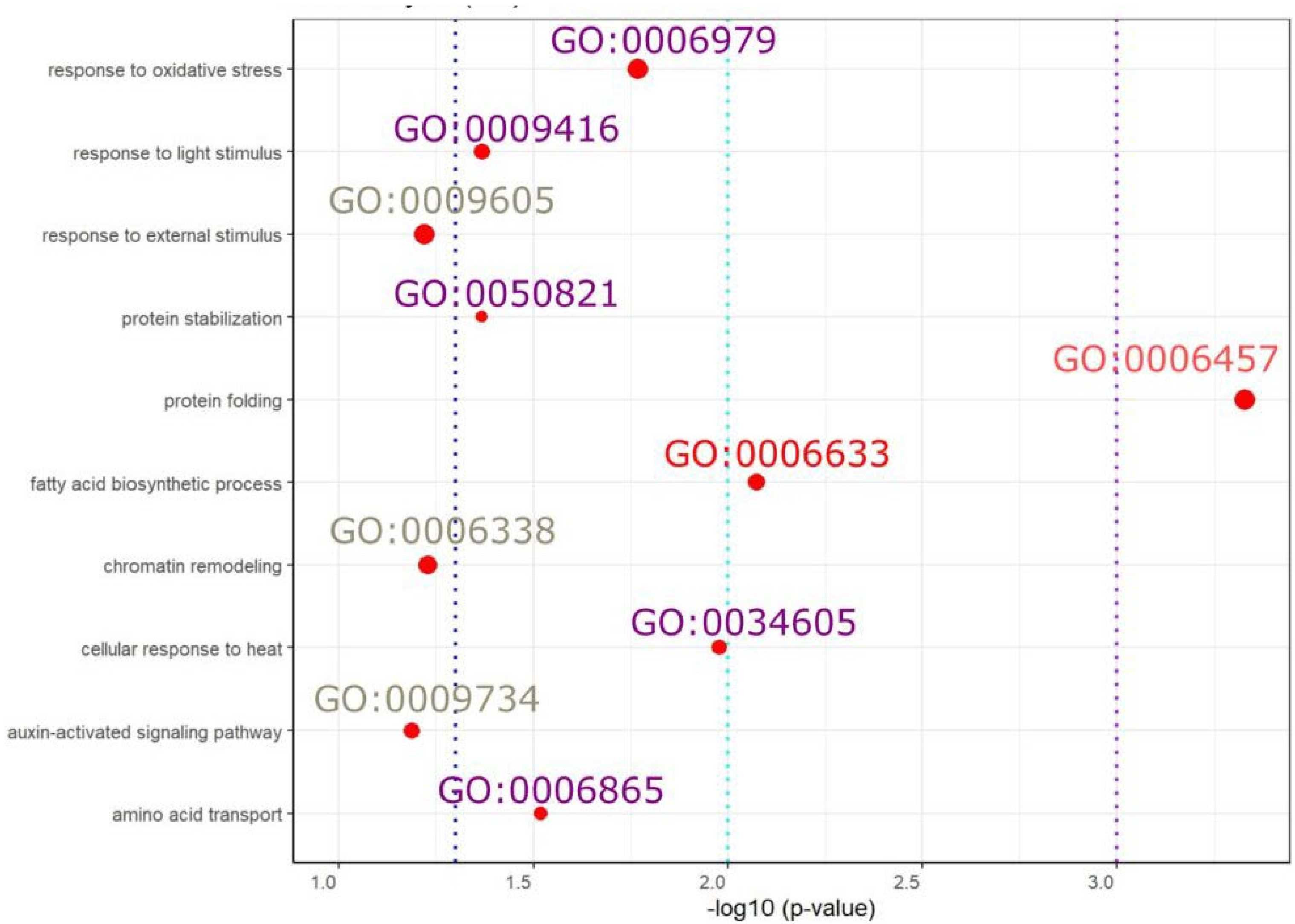

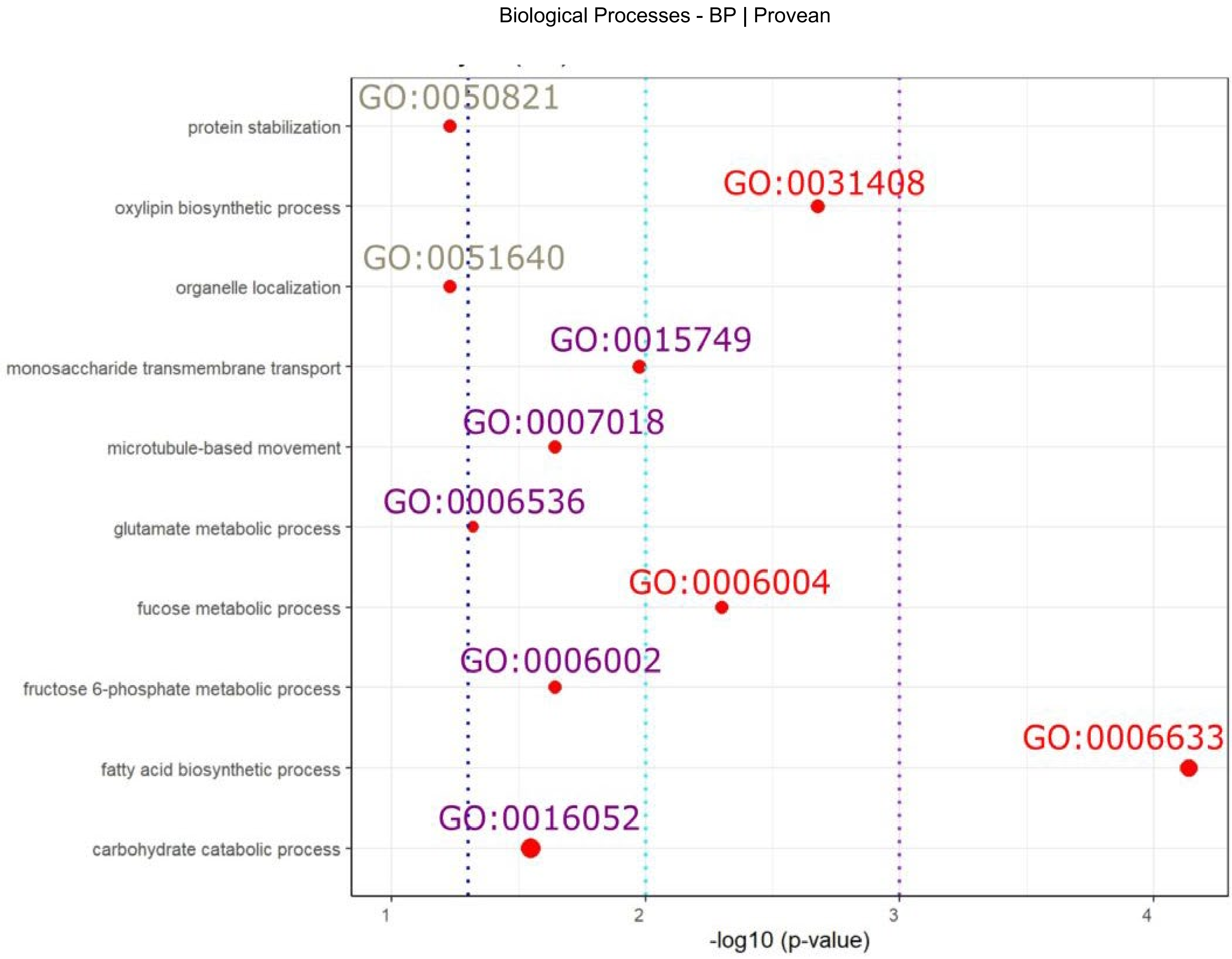

Biological Processes - BP | Provean
GO:0050821
protein stabilization
GO:0031408
oxylipin biosynthetic process
GO:0051640
organelle localization
GO:0015749
monosaccharide transmembrane transport
GO:0007018
microtubule-based movement
GO:0006536
glutamate metabolic process
GO:0006004
fucose metabolic process
GO:0006002
fructose 6-phosphate metabolic process
GO:0006633
fatty acid biosynthetic process
GO:0016052
carbohydrate catabolic process
1
2
3
4
-log10 (p-value)

B.

Biological Processes - BP | SnpEff

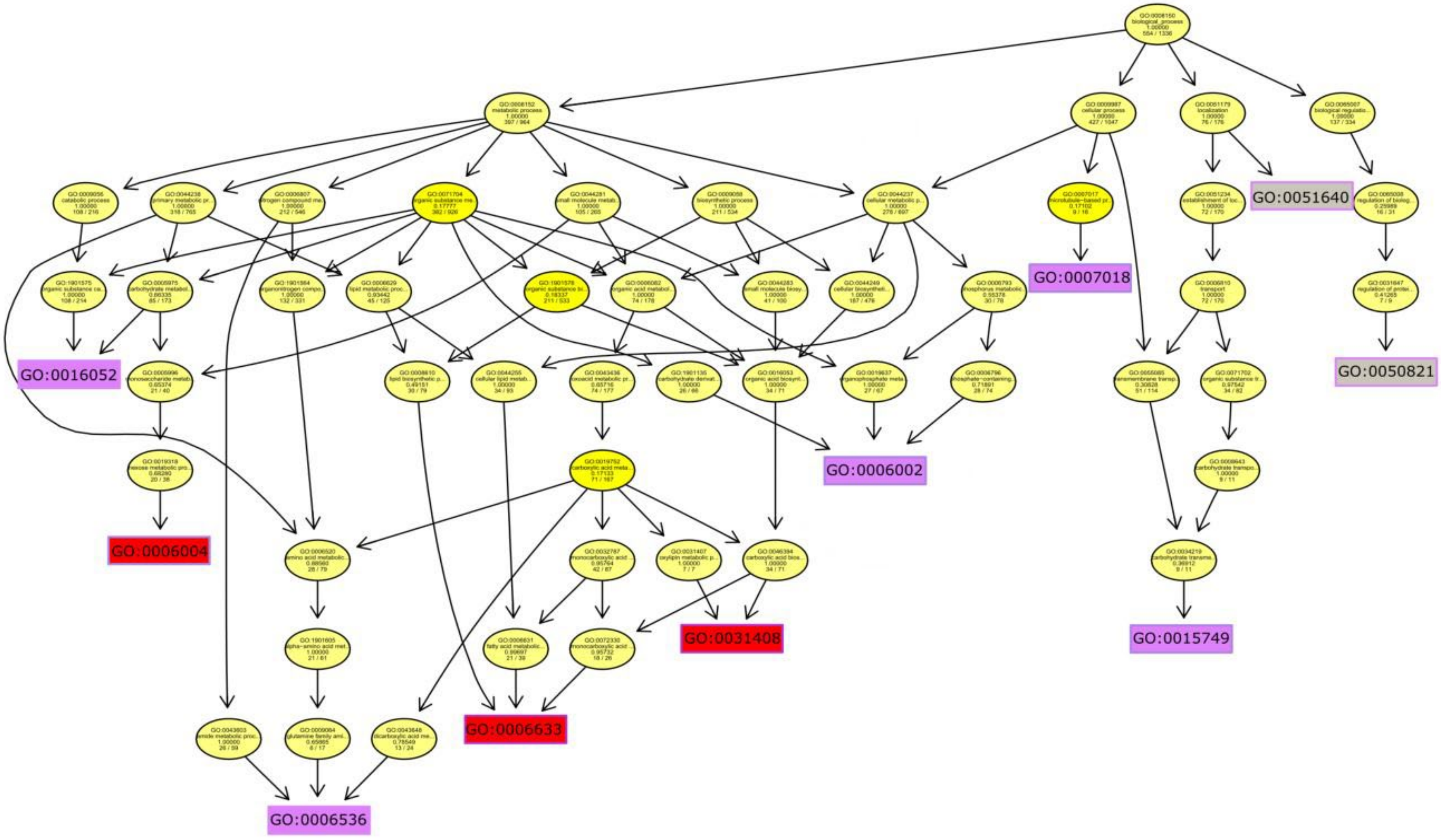

Biological Processes - BP | Provean

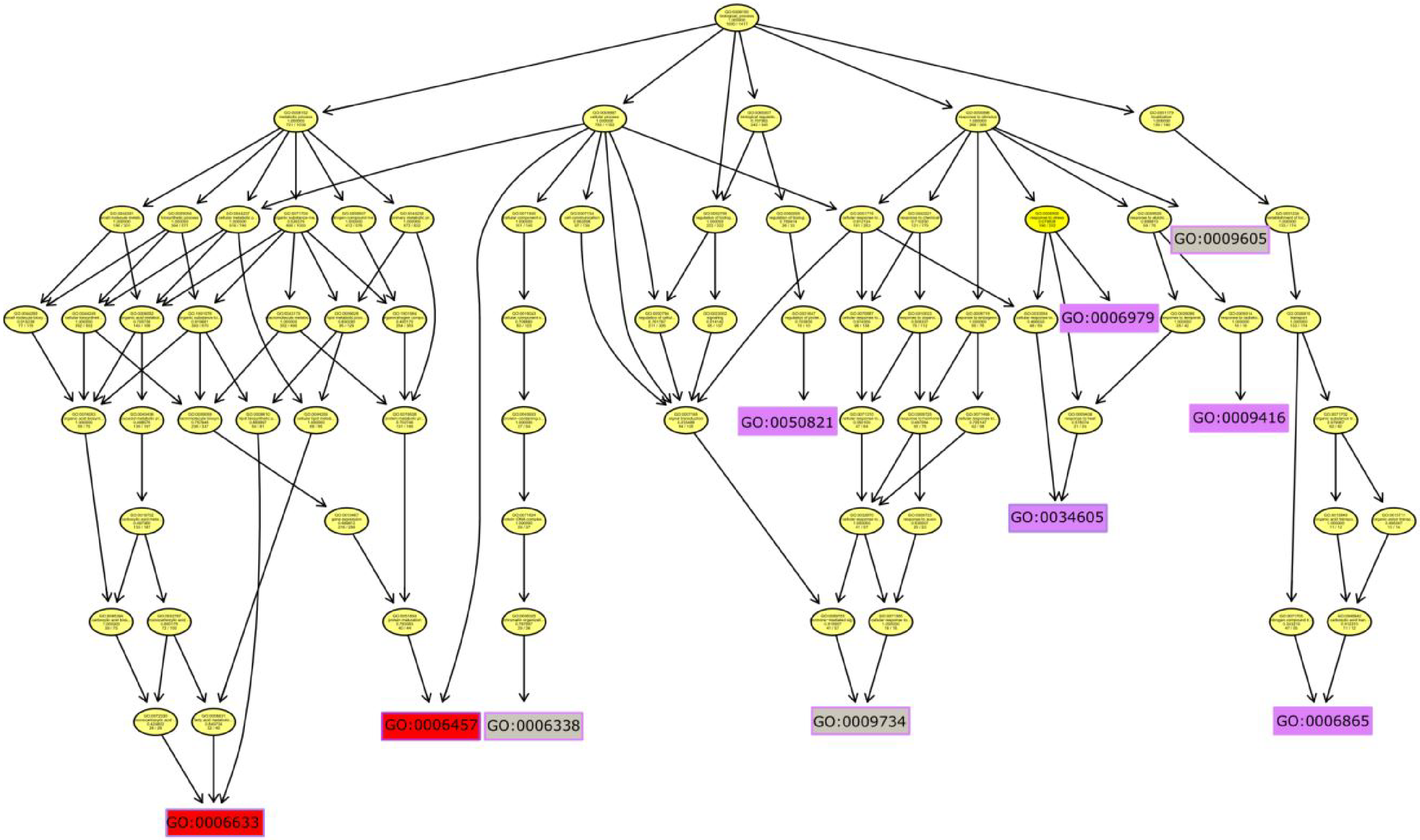

A.

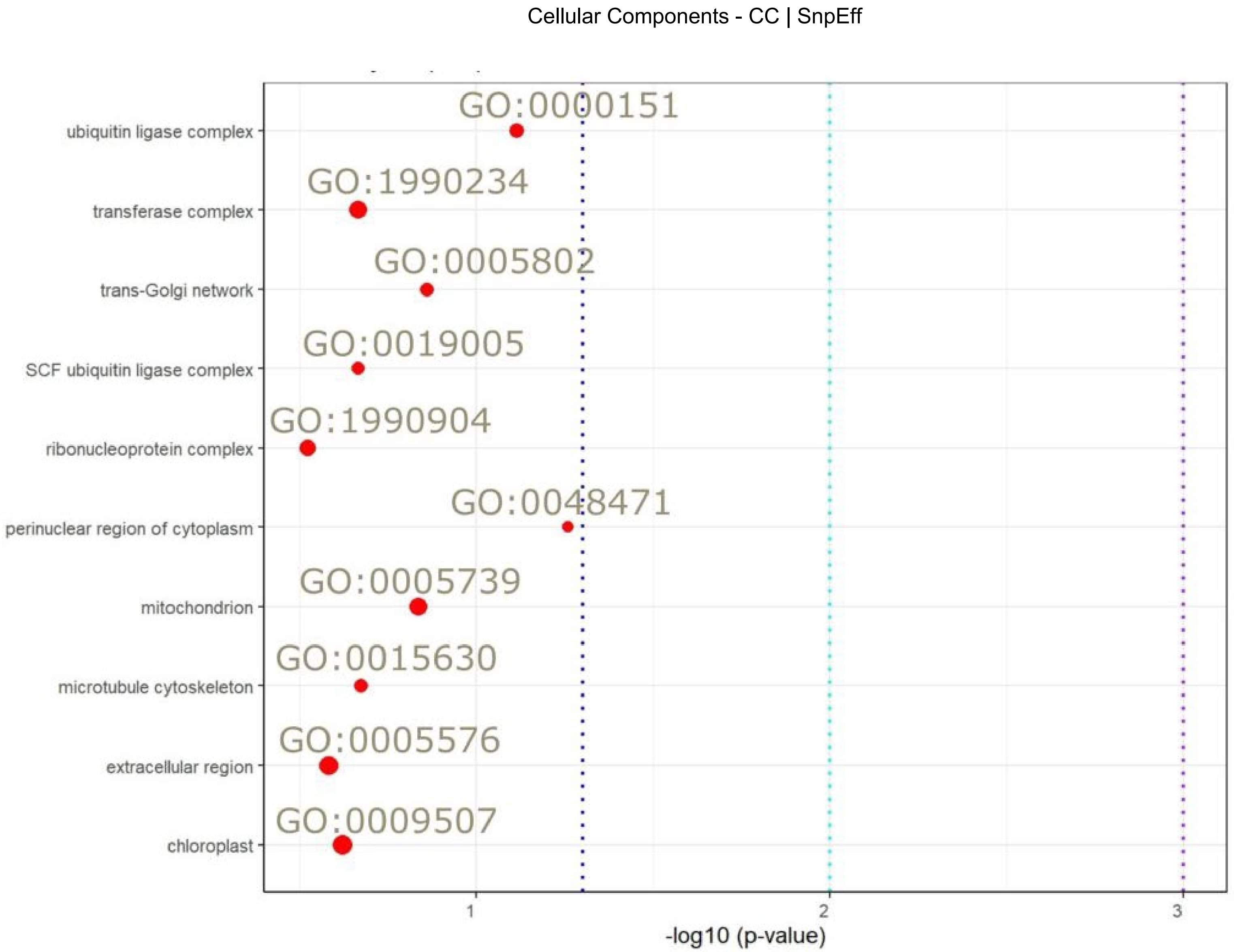
Cellular Components - CC | SnpEff
ubiquitin ligase complex
transferase complex
trans-Golgi network
SCF ubiquitin ligase complex
ribonucleoprotein complex
perinuclear region of cytoplasm
mitochondrion
microtubule cytoskeleton
extracellular region
chloroplast
GO:0000151
GO:1990234
GO:0005802
GO:0019005
GO:1990904
GO:0048471
GO:0005739
GO:0015630
GO:0005576
GO:0009507
1
2
3
-log10 (p-value)

Cellular Components - CC | Provean

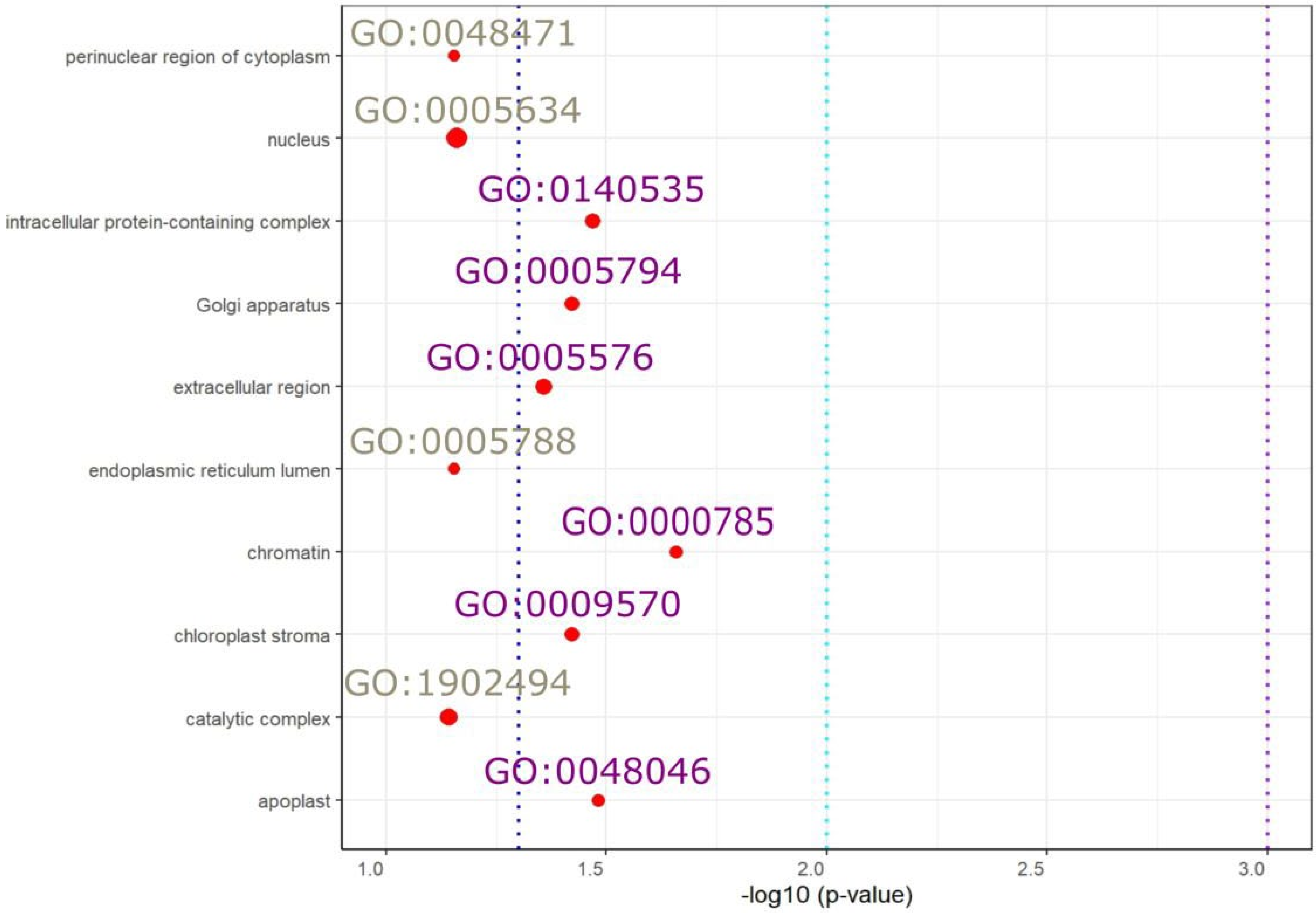

B.

Cellular Components - CC | SnpEff

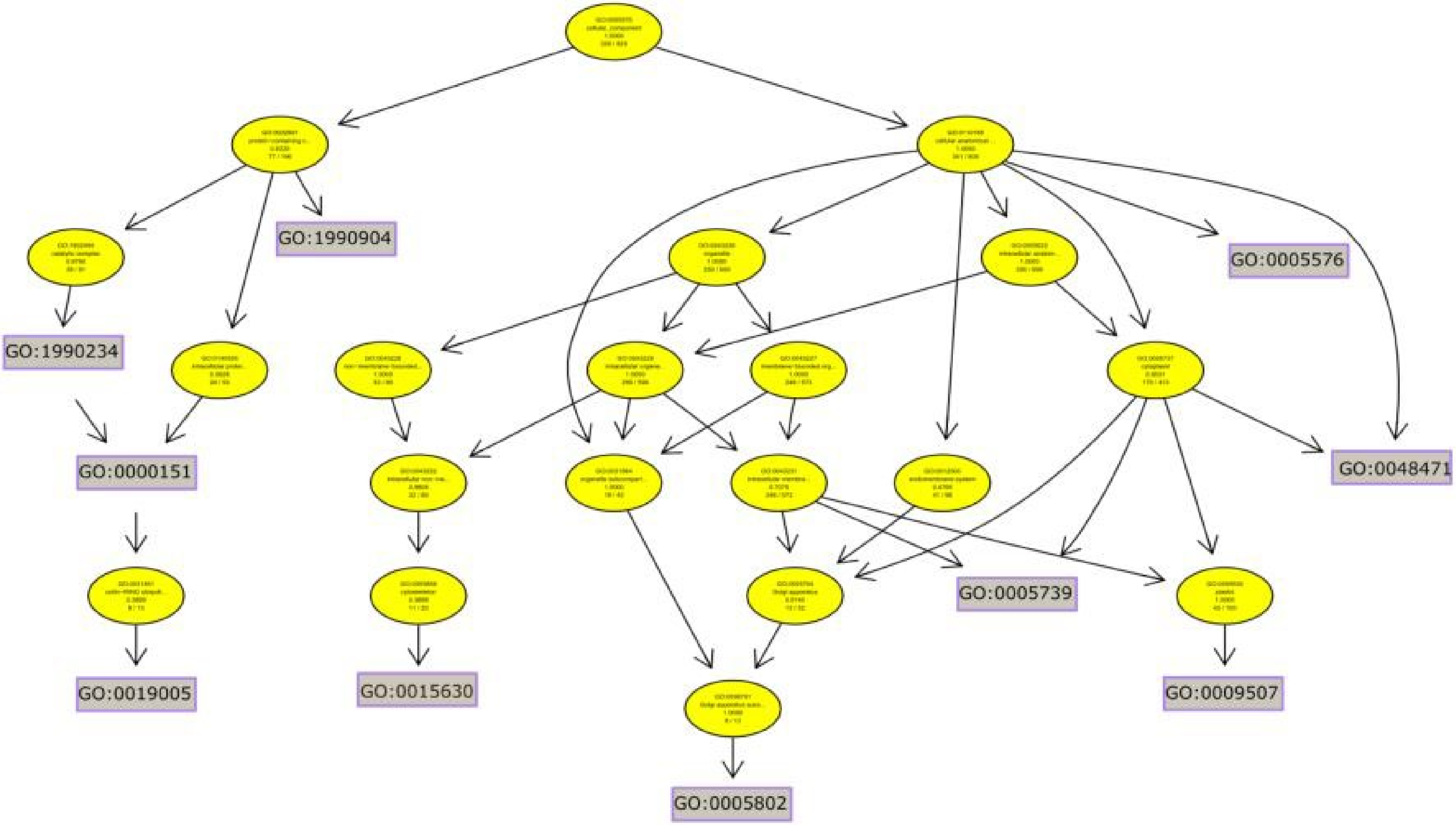

Cellular Components - CC | Provean

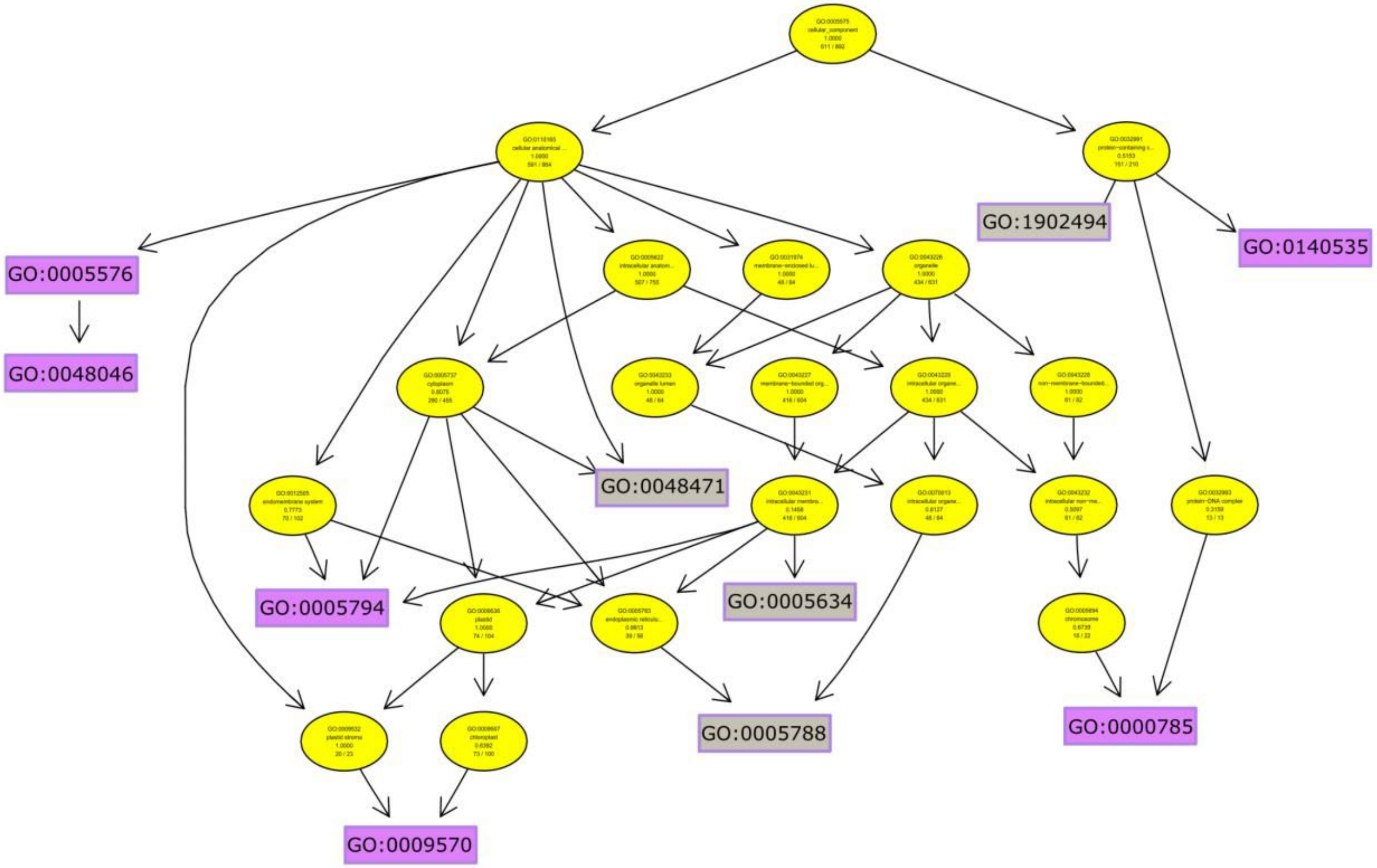

A.

Molecular Functions - MF | SnpEff

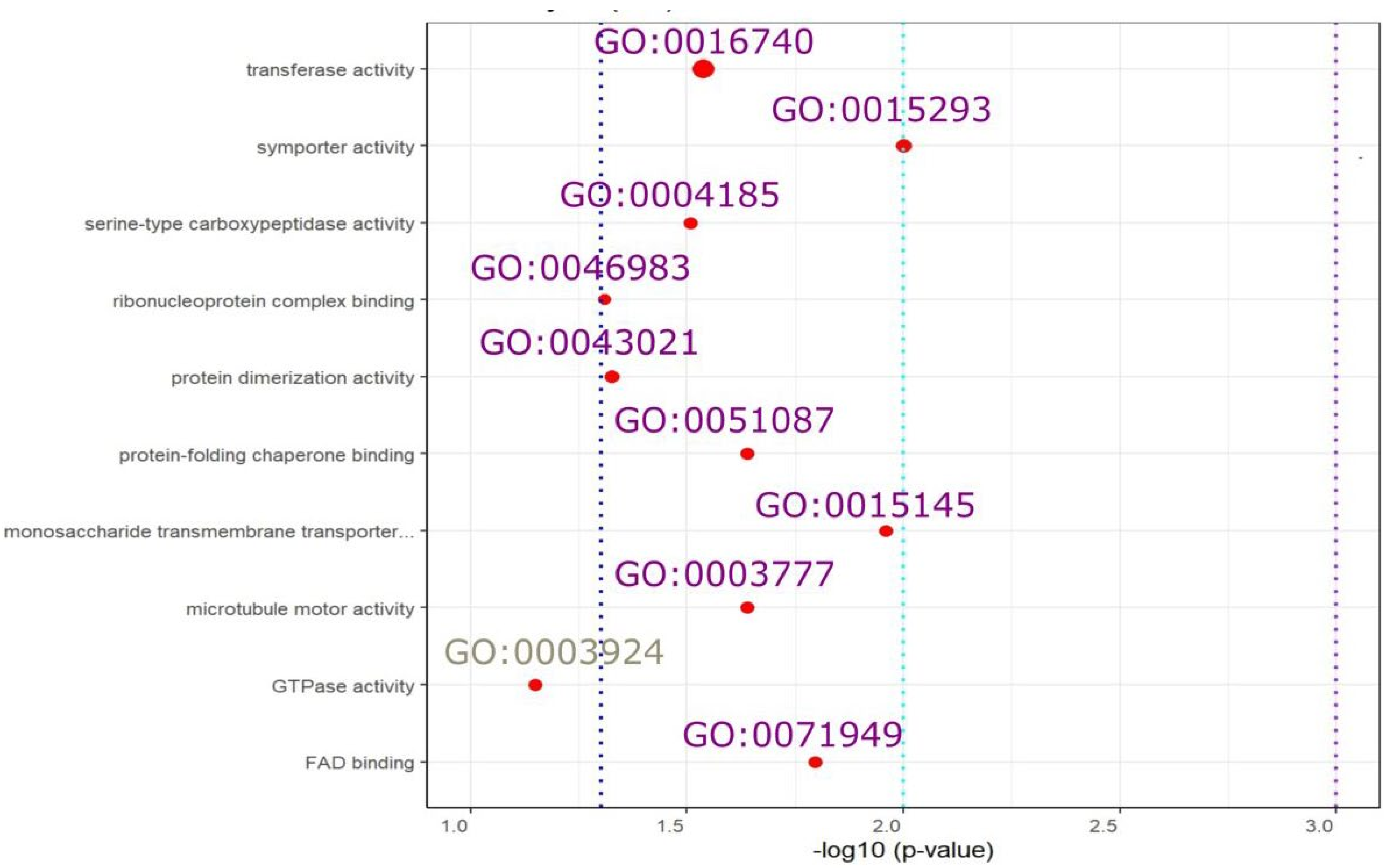

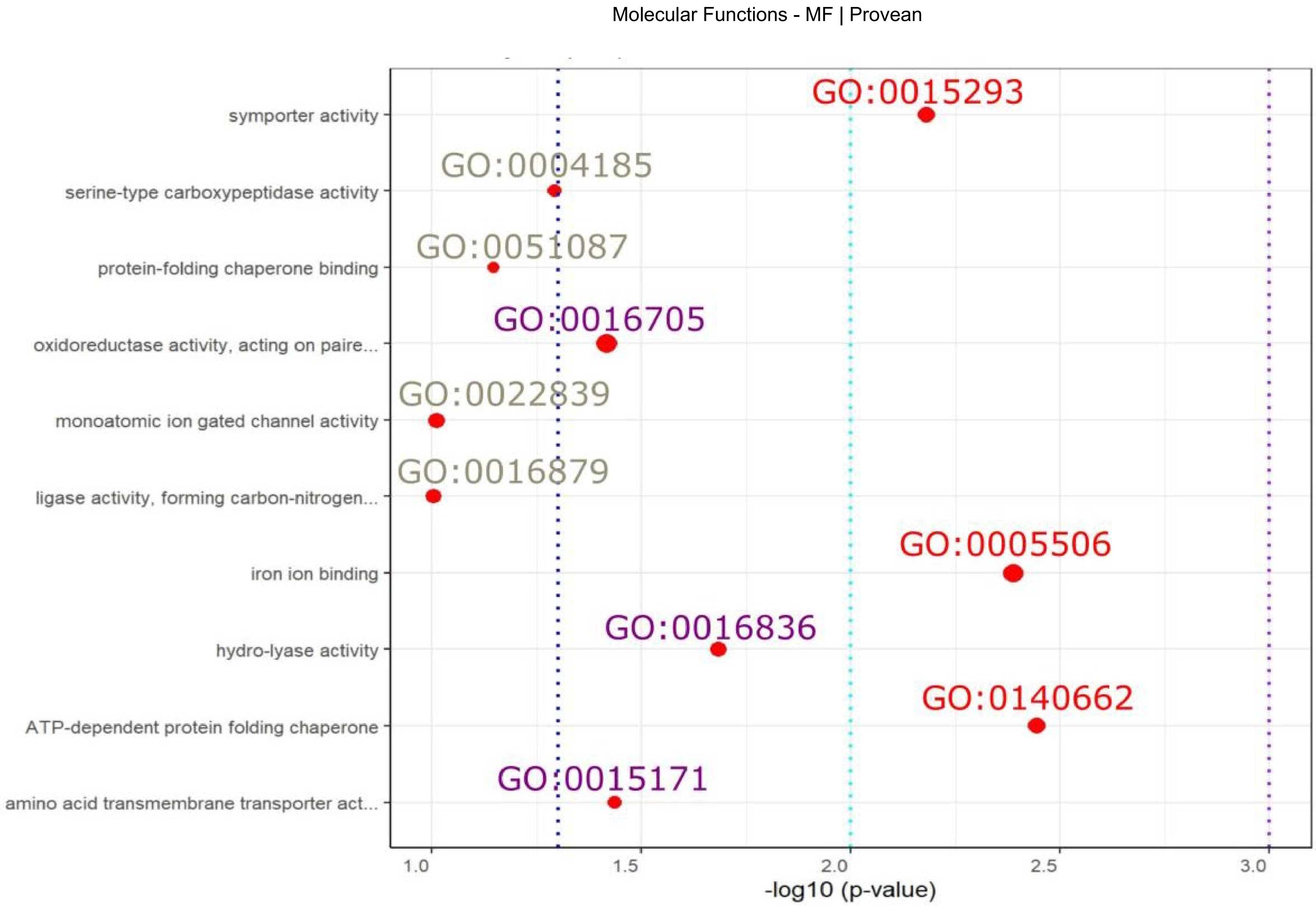

Molecular Functions - MF | Provean
symporter activity
serine-type carboxypeptidase activity
protein-folding chaperone binding
oxidoreductase activity, acting on paire...
monoatomic ion gated channel activity
ligase activity, forming carbon-nitrogen...
iron ion binding
hydro-lyase activity
ATP-dependent protein folding chaperone
amino acid transmembrane transporter act...
GO:0015293
GO:0004185
GO:0051087
GO:0016705
GO:0022839
GO:0016879
GO:0005506
GO:0016836
GO:0140662
GO:0015171
1.0
1.5
2.0
2.5
3.0
-log10 (p-value)

B.

Molecular Functions - MF | SnpEff

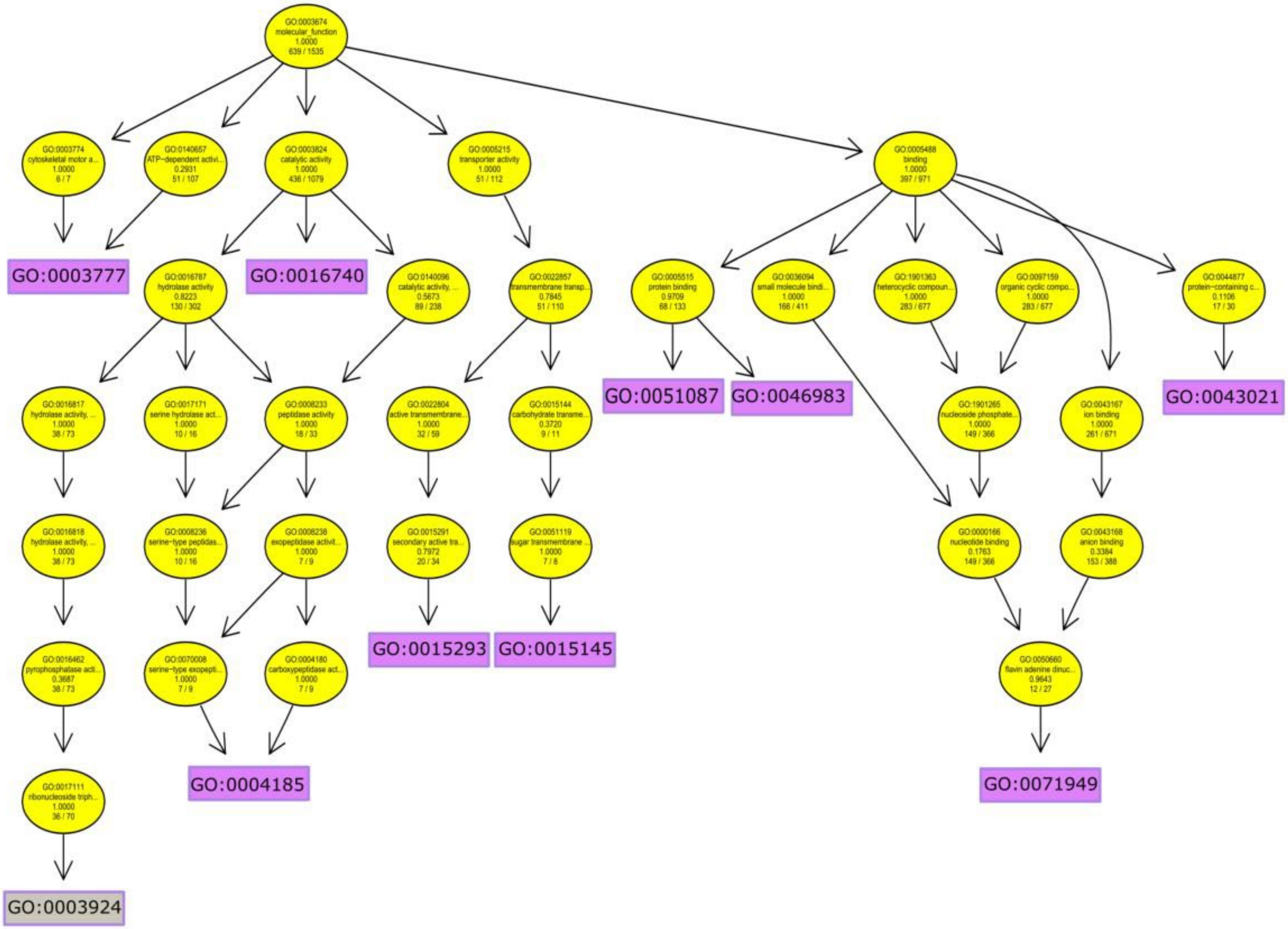

Molecular Functions - MF | Provean

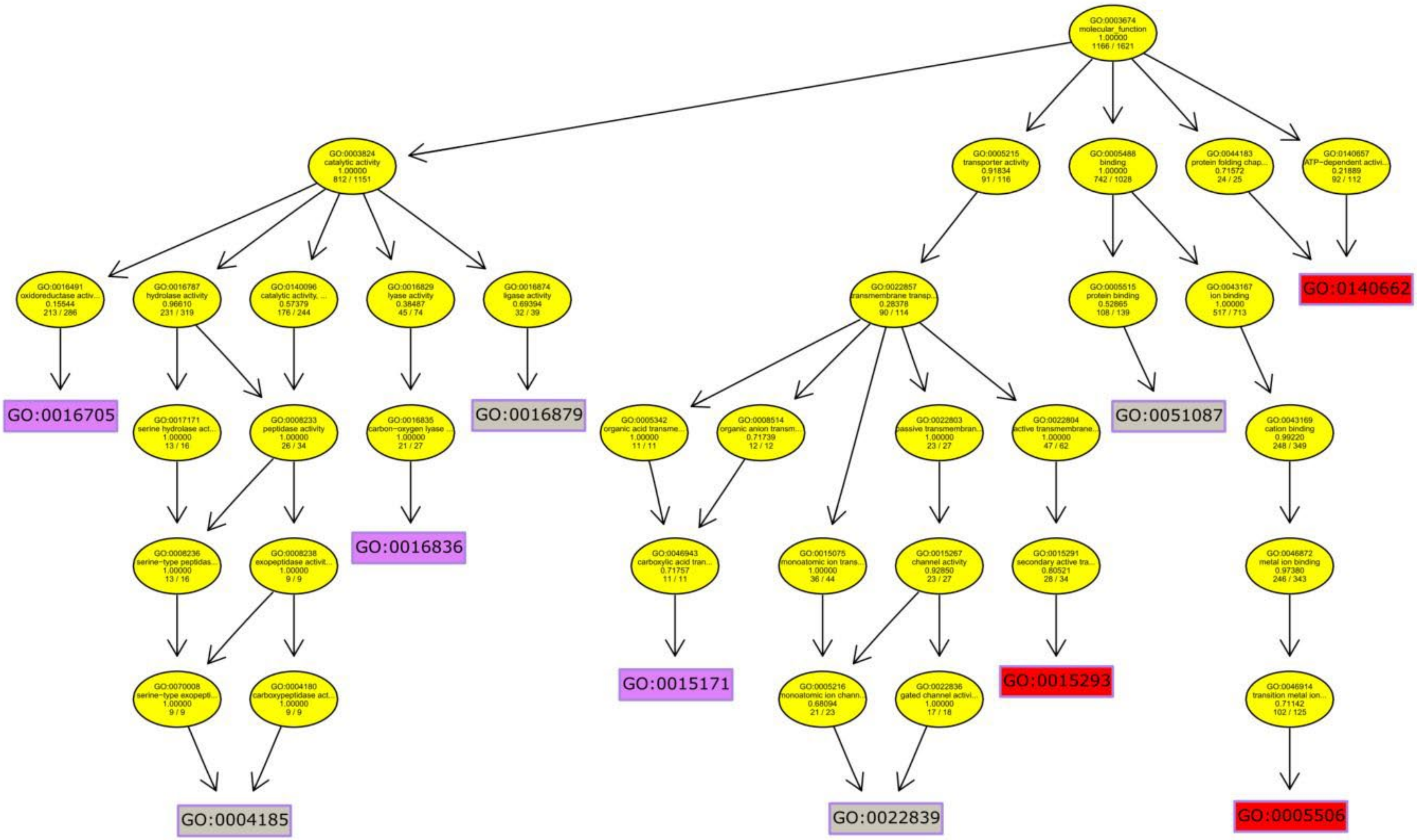

**Table S1**. Sampling strategy, locality information and estimates of genetic diversity and genetic load for the two Corsican local datasets (A) and the range-wide dataset of maritime pine (B).

A. *CORSICA-capture* and *CORSICA-array* datasets

| ***Locality Nr.*** | ***Locality Code*** | ***Country*** | ***Provenance*** | ***Lat.*** | ***Long.*** | ***Alt.*** | ***CORSICA capture dataset*** | | | | ***CORSICA array dataset*** | | |
|---|---|---|---|---|---|---|---|---|---|---|---|---|---|
| | | | | | | | ***N*** | ***Recessive GL*** | ***Additive GL*** | ***Ho*** | ***N*** | ***Recessive GL*** | ***Additive GL*** |
| 1 | VEN | France | Ventilegne | 41.44 | 9.12 | 50 | 24 | 0.749 | 0.767 | 0.047 | 16 | 0.468 | 0.495 |
| 2 | CAG | France | Cagna | 41.6 | 9.14 | 1040 | | | | | 12 | 0.430 | 0.478 |
| 3 | OSP | France | Ospedale | 41.65 | 9.2 | 800 | 2 | 0.746 | 0.772 | 0.055 | 9 | 0.421 | 0.481 |
| 4 | BAR | France | Barocaggio | 41.68 | 9.2 | 950 | | | | | 12 | 0.441 | 0.497 |
| 5 | BRU | France | Bruscaju | 41.72 | 9.3 | 300 | | | | | 12 | 0.419 | 0.483 |
| 6 | VAL | France | Vallemalla | 41.75 | 9.02 | 690 | 34 | 0.746 | 0.763 | 0.052 | | | |
| 7 | ZON | France | Zonza | 41.75 | 9.19 | 760 | 32 | 0.749 | 0.772 | 0.054 | | | |
| 8 | AUL | France | Aullene | 41.78 | 9.05 | 1105 | 27 | 0.751 | 0.767 | 0.051 | 16 | 0.433 | 0.482 |
| 9 | BAV | France | Bavella | 41.8 | 9.24 | 1020 | 33 | 0.743 | 0.760 | 0.052 | 16 | 0.447 | 0.486 |
| 10 | ARZ | France | Arza | 41.81 | 9.25 | 700 | 11 | 0.757 | 0.781 | 0.047 | | | |
| 11 | LAR | France | Larone | 41.83 | 9.27 | 590 | 9 | 0.759 | 0.780 | 0.050 | | | |
| 12 | CAL | France | Calzatoju | 41.83 | 9.32 | 130 | | | | | 12 | 0.425 | 0.466 |
| 13 | UTR | France | Utriolu | 41.85 | 9.32 | 405 | | | | | 12 | 0.431 | 0.484 |
| 14 | TOV | France | Tova | 41.88 | 9.3 | 975 | 12 | 0.758 | 0.780 | 0.048 | | | |
| 15 | PUN | France | Punta | 41.95 | 8.7 | 650 | | | | | 12 | 0.432 | 0.474 |
| 16 | PTA | France | Pineta | 41.97 | 9.04 | 750 | 31 | 0.748 | 0.767 | 0.049 | 16 | 0.471 | 0.495 |
| 17 | ANI | France | Ania | 41.97 | 9.28 | 630 | | | | | 12 | 0.453 | 0.491 |
| 18 | PIA | France | Pinia | 42 | 9.46 | 10 | 29 | 0.742 | 0.761 | 0.053 | 16 | 0.438 | 0.493 |
| 19 | VER | France | Vero | 42.07 | 8.93 | 575 | 29 | 0.742 | 0.761 | 0.051 | 16 | 0.447 | 0.488 |
| 20 | OME | France | Omenino | 42.13 | 9.14 | 1080 | | | | | 12 | 0.419 | 0.471 |
| 21 | SOR | France | Sorba | 42.14 | 9.19 | 1140 | 29 | 0.749 | 0.772 | 0.051 | 16 | 0.451 | 0.482 |
| 22 | PAS | France | Pastricciola | 42.15 | 9 | 700 | 34 | 0.735 | 0.759 | 0.051 | 16 | 0.429 | 0.479 |
| 23 | GUA | France | Guagno | 42.17 | 8.88 | 500 | | | | | 12 | 0.437 | 0.481 |
| 24 | CER | France | Cervello | 42.2 | 9.13 | 795 | | | | | 12 | 0.430 | 0.491 |

| | | | | | | | | | | | | | |
|---|---|---|---|---|---|---|---|---|---|---|---|---|---|
| 25 | RES | France | Restonica | 42.27 | 9.1 | 700 | 29 | 0.755 | 0.776 | 0.054 | 16 | 0.434 | 0.489 |
| 26 | OMA | France | Ominanda | 42.33 | 9.12 | 890 | | | | | 12 | 0.457 | 0.491 |
| 27 | PER | France | Perticato | 42.36 | 8.79 | 360 | | | | | 10 | 0.450 | 0.516 |
| 28 | PIN | France | Pineto | 42.43 | 9.23 | 365 | | | | | 12 | 0.451 | 0.484 |
| 29 | BON | France | Bonifatu | 42.45 | 8.83 | 440 | 28 | 0.752 | 0.772 | 0.053 | 16 | 0.449 | 0.480 |
| 30 | TAR | France | Tartagine | 42.49 | 8.97 | 960 | | | | | 12 | 0.457 | 0.496 |

B. *GLOBAL-array* dataset

| ***Locality Nr.*** | ***Locality Code*** | ***Country*** | ***Provenance*** | **Genepool** | ***Lat.*** | ***Long.*** | ***Alt.*** | ***GLOBAL-array* dataset** | | |
|---|---|---|---|---|---|---|---|---|---|---|
| | | | | | | | | **N** | **Recessive GL** | **Additive GL** |
| 1 | PLE | France | Pleucadec | Atlantic France | 47.78 | -2.34 | 70 | 17 | 0.467 | 0.519 |
| 2 | STJ | France | St-Jean des Monts | Atlantic France | 46.76 | -2.03 | 6 | 25 | 0.466 | 0.517 |
| 3 | OLO | France | Olonne sur Mer | Atlantic France | 46.57 | -1.83 | 12 | 24 | 0.459 | 0.510 |
| 4 | VER | France | Le Verdon | Atlantic France | 45.55 | -1.09 | 10 | 25 | 0.447 | 0.499 |
| 5 | HOU | France | Hourtin | Atlantic France | 45.18 | -1.15 | 28 | 25 | 0.463 | 0.511 |
| 6 | MIM | France | Mimizan | Atlantic France | 44.13 | -1.3 | 18 | 17 | 0.457 | 0.500 |
| 7 | PET | France | Petrocq | Atlantic France | 44.06 | -1.3 | 21 | 22 | 0.465 | 0.508 |
| 8 | LAM | Spain | Lamuño | Iberian Atlantic | 43.56 | -6.22 | 119 | 9 | 0.452 | 0.470 |
| 9 | PUE | Spain | Puerto de Vega | Iberian Atlantic | 43.55 | -6.63 | 81 | 7 | 0.463 | 0.493 |
| 10 | CAD | Spain | Cadavedo | Iberian Atlantic | 43.54 | -6.42 | 164 | 8 | 0.443 | 0.481 |
| 11 | SIE | Spain | Sierra de Barcia | Iberian Atlantic | 43.53 | -6.49 | 264 | 8 | 0.422 | 0.491 |
| 12 | CAS | Spain | Castropol | Iberian Atlantic | 43.5 | -6.98 | 158 | 8 | 0.478 | 0.494 |
| 13 | ARM | Spain | Armayán | Iberian Atlantic | 43.3 | -6.46 | 559 | 7 | 0.496 | 0.486 |
| 14 | ALT | Spain | Alto de la Llama | Iberian Atlantic | 43.28 | -6.49 | 518 | 8 | 0.480 | 0.491 |
| 15 | SEG | Spain | Sergude (Huerto Semillero) | Iberian Atlantic | 42.82 | -8.45 | 309 | 19 | 0.469 | 0.494 |
| 16 | SAC | Spain | As Neves, San Cipriano de Ribarteme | Iberian Atlantic | 42.12 | -8.37 | 415 | 8 | 0.421 | 0.471 |
| 17 | LEI | Portugal | Leiria | Iberian Atlantic | 39.78 | -8.96 | 79 | 21 | 0.460 | 0.492 |
| 18 | SAL | Spain | San Leonardo de Yagüe | Central Spain | 41.83 | -3.06 | 1077 | 10 | 0.454 | 0.516 |
| 19 | BAY | Spain | Bayubas de Abajo | Central Spain | 41.52 | -2.88 | 925 | 15 | 0.465 | 0.507 |
| 20 | CUE | Spain | Cuéllar | Central Spain | 41.33 | -4.25 | 754 | 23 | 0.462 | 0.513 |
| 21 | COC | Spain | Coca | Central Spain | 41.25 | -4.5 | 784 | 15 | 0.484 | 0.515 |
| 22 | CAR | Spain | Carbonero el Mayor | Central Spain | 41.17 | -4.28 | 844 | 6 | 0.544 | 0.531 |
| 23 | VAL | Spain | Valdemaqueda | Central Spain | 40.51 | -4.31 | 970 | 10 | 0.457 | 0.496 |
| 24 | CEN | Spain | Cenicientos | Central Spain | 40.28 | -4.49 | 1079 | 9 | 0.480 | 0.504 |

| | | | | | | | | | | |
|---|---|---|---|---|---|---|---|---|---|---|
| 25 | ARN | Spain | Arenas de San Pedro | Central Spain | 40.19 | -5.12 | 663 | 14 | 0.492 | 0.523 |
| 26 | OLB | Spain | Olba | Central Spain | 40.17 | -0.62 | 989 | 19 | 0.453 | 0.507 |
| 27 | QUA | Spain | Quatretonda | Southern Spain | 38.97 | -0.36 | 459 | 16 | 0.466 | 0.509 |
| 28 | BON | Spain | Boniches | Central Spain | 39.98 | -1.66 | 1098 | 8 | 0.508 | 0.521 |
| 29 | ORI | Spain | Oria | Southern Spain | 37.51 | -2.32 | 1231 | 22 | 0.483 | 0.512 |
| 30 | COM | Spain | Competa | Southern Spain | 36.83 | -3.93 | 904 | 3 | 0.532 | 0.509 |
| 31 | MAD | Morocco | Madisouka | Morocco | 35.18 | -5.23 | 1302 | 1 | 0.520 | 0.531 |
| 32 | TAM | Morocco | Tamrabta | Morocco | 33.6 | -5.02 | 1729 | 12 | 0.451 | 0.502 |
| 33 | PIA | France | Pinia | Corsica | 42.02 | 9.46 | 8 | 13 | 0.502 | 0.529 |
| 34 | PIE | France | Pineta | Corsica | 41.97 | 9.04 | 909 | 10 | 0.486 | 0.516 |

**Table S2**. Linear mixed models of the genetic load based on SnpEff annotations with phenotypic traits related to water-use efficiency, tree growth and bud-burst in maritime pine for the two regional datasets from Corsica, including or not altitude of the provenance as random effect: (A) *CORSICA-capture* and (B) *CORSICA-array,* as well as for (C) the global dataset *GLOBAL-array*. For the latter dataset, correlation with survival was also tested. The recessive genetic load (GL) focusses on the number of homozygous sites containing deleterious mutations whereas the additive genetic load indicates the total number of deleterious alleles in homozygous and heterozygous sites. Traits used were: carbon isotope discrimination ($\delta^{13}C$) for water-use efficiency, tree height in cm for tree growth, and Julian day of entry in accumulated degree-days from the first day of the year for bud-burst. CI, confidence intervals; *p*, *p*-value based on conditional F-tests with Kenward-Roger approximation for the degrees of freedom; $\sigma^2$, residual variance; $\tau_{00}$, random effect variances; ICC, intraclass correlation coefficient; N, number of random effects groups; Marg. $R^2$, marginal goodness of fit (fixed effects); Cond. $R^2$, conditional goodness of fit (fixed and random effects). For the recessive GL tests in Corsica, bud-burst exhibited no variation with altitude and block in the capture and array datasets, respectively.

A. *CORSICA-capture* dataset

| | | $\delta^{13}C$ | | | $\delta^{13}C$ | |
|---|---|---|---|---|---|---|
| *Predictors* | *Estimates* | *CI* | *p* | *Estimates* | *CI* | *p* |
| (Intercept) | -34.81 | -41.54 – -28.08 | **<0.001** | -34.22 | -40.89 – -27.54 | **<0.001** |
| recessive GL | 8.00 | -1.00 – 17.01 | 0.081 | | | |
| additive GL | | | | 7.03 | -1.67 – 15.72 | 0.112 |
| **Random Effects** | | | | | | |
| $\sigma^2$ | 0.45 | | | 0.45 | | |
| $\tau_{00}$ | 0.06 block:plot_id | | | 0.06 block:plot_id | | |
| | 0.05 plot_id | | | 0.06 plot_id | | |
| ICC | 0.20 | | | 0.20 | | |
| N | 3 block | | | 3 block | | |
| | 3 plot_id | | | 3 plot_id | | |
| Observations | 93 | | | 93 | | |
| Marg / Cond $R^2$ | 0.030 / 0.221 | | | 0.025 / 0.223 | | |

| | height_cm | | | height_cm | | |
|---|---|---|---|---|---|---|
| *Predictors* | *Estimates* | *CI* | *p* | *Estimates* | *CI* | *p* |
| (Intercept) | 521.04 | 231.11 – 810.97 | **0.001** | 432.89 | 133.93 – 731.85 | **0.005** |
| Recessive GL | -451.22 | -837.59 – -64.84 | **0.022** | | | |
| Additive GL | | | | -324.58 | -712.26 – 63.09 | 0.100 |
| **Random Effects** | | | | | | |
| $\sigma^2$ | 1571.70 | | | 1596.33 | | |
| $\tau_{00}$ | 256.50 altitude | | | 260.84 altitude | | |
| | 142.91 block | | | 142.24 block | | |
| ICC | 0.20 | | | 0.20 | | |
| N | 5 block | | | 5 block | | |
| | 14 altitude | | | 14 altitude | | |
| Observations | 168 | | | 168 | | |
| Mar/ Cond $R^2$ | 0.027 / 0.224 | | | 0.015 / 0.213 | | |

| | height_cm | | | height_cm | | |
|---|---|---|---|---|---|---|
| *Predictors* | *Estimates* | *CI* | *p* | *Estimates* | *CI* | *p* |
| (Intercept) | 538.59 | 241.24 – 835.93 | **<0.001** | 439.35 | 134.96 – 743.74 | **0.005** |
| recessive GL | -477.96 | -874.01 – -81.92 | **0.018** | | | |
| additive GL | | | | -324.58 | -712.26 – 63.09 | 0.100 |
| **Random Effects** | | | | | | |
| $\sigma^2$ | 1745.53 | | | 1776.64 | | |
| $\tau_{00}$ | 295.30 $_{block}$ | | | 285.86 $_{block}$ | | |
| ICC | 0.14 | | | 0.14 | | |
| N | 5 $_{block}$ | | | 5 $_{block}$ | | |
| Observations | 168 | | | 168 | | |
| Marg / Cond $R^2$ | 0.029 / 0.170 | | | 0.015 / 0.152 | | |

| | bud-burst | | | bud-burst | | |
|---|---|---|---|---|---|---|
| *Predictors* | *Estimates* | *CI* | *p* | *Estimates* | *CI* | *p* |
| (Intercept) | 2995.69 | 2175.28 – 3816.09 | **<0.001** | 3097.23 | 2271.58 –3922.89 | **<0.001** |
| recessive GL | -1325.76 | -2421.16 – -230.36 | **0.018** | | | |
| additive GL | | | | -1424.13 | -2498.37 –-349.89 | **0.010** |
| **Random Effects** | | | | | | |
| $\sigma^2$ | 9854.70 | | | 9778.63 | | |
| $\tau_{00}$ | 1699.19 $_{plot\_id}$ | | | 1643.37 $_{plot\_id}$ | | |
| ICC | 0.15 | | | 0.14 | | |
| N | 3 $_{plot\_id}$ | | | 3 $_{plot\_id}$ | | |
| Observations | 139 | | | 139 | | |
| Marg / Cond $R^2$ | 0.037 / 0.178 | | | 0.044 / 0.182 | | |

| | bud-burst | | | bud-burst | | |
|---|---|---|---|---|---|---|
| *Predictors* | *Estimates* | *CI* | *p* | *Estimates* | *CI* | *p* |
| (Intercept) | 3149.47 | 2343.98 – 3954.95 | **<0.001** | 3255.17 | 2447.97 – 4062.37 | **<0.001** |
| Recessive GL | -1529.78 | -2604.18 – -455.38 | **0.006** | | | |
| Additive GL | | | | -1628.95 | -2678.64 – -579.27 | **0.003** |
| **Random Effects** | | | | | | |
| $\sigma^2$ | 9047.80 | | | 8941.76 | | |
| $\tau_{00}$ | 1076.31 $_{altitude}$ | | | 1104.54 $_{altitude}$ | | |
| | 1506.41 $_{plot\_id}$ | | | 1517.54 $_{plot\_id}$ | | |
| ICC | 0.22 | | | 0.23 | | |
| N | 3 $_{plot\_id}$ | | | 3 $_{plot\_id}$ | | |
| | 11 $_{altitude}$ | | | 11 $_{altitude}$ | | |
| Observations | 139 | | | 139 | | |
| Marg / Cond $R^2$ | 0.048 / 0.260 | | | 0.056 / 0.270 | | |

B. *CORSICA-array* dataset

| | δ13C | | | δ13C | | |
|---|---|---|---|---|---|---|
| *Predictors* | *Estimates* | *CI* | *p* | *Estimates* | *CI* | *p* |
| (Intercept) | -28.49 | -29.69 – -27.29 | **<0.001** | -27.94 | -29.82 – -26.06 | **<0.001** |
| Recessive GL | -1.05 | -3.49 – 1.40 | 0.398 | | | |
| Additive GL | | | | -1.96 | -5.63 – 1.70 | 0.290 |
| **Random Effects** | | | | | | |
| $\sigma^2$ | 0.34 | | | 0.34 | | |
| $\tau_{00}$ | 0.17 $_{block:plot\_id}$ | | | 0.26 $_{plot\_id}$ | | |
| | 0.00 $_{plot\_id}$ | | | | | |
| ICC | 0.34 | | | 0.43 | | |
| N | 3 $_{block}$ | | | 3 $_{plot\_id}$ | | |
| | 3 $_{plot\_id}$ | | | | | |
| Observations | 120 | | | 120 | | |
| Marg. / Cond. $R^2$ | 0.004 / 0.344 | | | 0.005 / 0.432 | | |

| | height_cm | | | height_cm | | |
|---|---|---|---|---|---|---|
| *Predictors* | *Estimates* | *CI* | *p* | *Estimates* | *CI* | *p* |
| (Intercept) | 263.83 | 216.70 – 310.96 | **<0.001** | 306.18 | 224.72 – 387.64 | **<0.001** |
| recessive GL | -162.09 | -263.97 – -60.21 | **0.002** | | | |
| additive GL | | | | -234.42 | -399.59 – -69.25 | **0.006** |
| **Random Effects** | | | | | | |
| $\sigma^2$ | 1868.37 | | | 1880.68 | | |
| $\tau_{00}$ | 140.44 $_{block:plot\_id}$ | | | 120.64 $_{block:plot\_id}$ | | |
| | 11.40 $_{plot\_id}$ | | | 24.58 $_{plot\_id}$ | | |
| ICC | 0.08 | | | 0.07 | | |
| N | 3 $_{block}$ | | | 3 $_{block}$ | | |
| | 3 $_{plot\_id}$ | | | 3 $_{plot\_id}$ | | |
| Observations | 334 | | | 334 | | |
| Marg / Cond $R^2$ | 0.027 / 0.100 | | | 0.022 / 0.092 | | |

| | height_cm | | | height_cm | | |
|---|---|---|---|---|---|---|
| Predictors | Estimates | CI | p | Estimates | CI | p |
| (Intercept) | 261.39 | 215.15 – 307.64 | **<0.001** | 309.95 | 229.95 – 389.95 | **<0.001** |
| recessive GL | -159.75 | -260.30 – -59.21 | **0.002** | | | |
| additive GL | | | | -244.72 | -407.07 – -82.38 | **0.003** |
| **Random Effects** | | | | | | |
| $\sigma^2$ | 1771.51 | | | 1772.73 | | |
| $\tau_{00}$ | 114.04 altitude | | | 128.37 altitude | | |
| | 84.66 block:plot_id | | | 64.68 block:plot_id | | |
| | 42.82 plot_id | | | 52.66 plot_id | | |
| ICC | 0.12 | | | 0.12 | | |
| N | 3 block | | | 3 block | | |
| | 3 plot_id | | | 3 plot_id | | |
| | 22 altitude | | | 22 altitude | | |
| Observations | 334 | | | 334 | | |
| Marg / Cond $R^2$ | 0.026 / 0.143 | | | 0.024 / 0.143 | | |

| | bud-burst | | | bud-burst | | |
|---|---|---|---|---|---|---|
| *Predictors* | *Estimates* | *CI* | *p* | *Estimates* | *CI* | *p* |
| (Intercept) | 1880.23 | 1712.71 – 2047.75 | **<0.001** | 1966.34 | 1678.36 – 2254.31 | **<0.001** |
| recessive GL | 275.86 | -87.30 – 639.02 | 0.135 | | | |
| additive GL | | | | 75.99 | -509.18 – 661.15 | 0.798 |
| **Random Effects** | | | | | | |
| $\sigma^2$ | 10728.44 | | | 10864.53 | | |
| $\tau_{00}$ | 30.54 $_{block:plot\_id}$ | | | 81.44 $_{block:plot\_id}$ | | |
| | 875.24 $_{plot\_id}$ | | | 934.26 $_{plot\_id}$ | | |
| ICC | 0.08 | | | 0.09 | | |
| N | 3 $_{block}$ | | | 3 $_{block}$ | | |
| | 3 $_{plot\_id}$ | | | 3 $_{plot\_id}$ | | |
| Observations | 153 | | | 153 | | |
| Marg / Cond $R^2$ | 0.014 / 0.091 | | | 0.000 / 0.086 | | |

| | bud-burst | | | bud-burst | | |
|---|---|---|---|---|---|---|
| *Predictors* | *Estimates* | *CI* | *p* | *Estimates* | *CI* | *p* |
| (Intercept) | 1876.19 | 1707.51 – 2044.87 | **<0.001** | 1958.86 | 1669.60 – 2248.13 | **<0.001** |
| recessive GL | 287.57 | -78.18 – 653.31 | 0.122 | | | |
| additive GL | | | | 93.92 | -494.39 – 682.22 | 0.753 |
| **Random Effects** | | | | | | |
| $\sigma^2$ | 10548.13 | | | 10726.39 | | |
| $\tau_{00}$ | 282.97 $_{altitude}$ | | | 231.53 $_{altitude}$ | | |
| | 849.75 $_{plot\_id}$ | | | 16.16 $_{block:plot\_id}$ | | |
| | | | | 937.30 $_{plot\_id}$ | | |
| ICC | 0.10 | | | 0.10 | | |
| N | 3 $_{plot\_id}$ | | | 3 $_{block}$ | | |
| | 8 $_{altitude}$ | | | 3 $_{plot\_id}$ | | |
| | | | | 8 $_{altitude}$ | | |
| Observations | 153 | | | 153 | | |
| Marg / Cond $R^2$ | 0.015 / 0.110 | | | 0.001 / 0.100 | | |

C. *GLOBAL-array* dataset

| | $\delta^{13}C$ | | | $\delta^{13}C$ | | |
|---|---|---|---|---|---|---|
| *Predictors* | *Estimates* | *CI* | *p* | *Estimates* | *CI* | *p* |
| (Intercept) | -26.40 | -27.27 – -25.54 | **<0.001** | -25.88 | -26.97 – -24.80 | **<0.001** |
| recessive GL | -0.38 | -1.32 – 0.56 | 0.427 | | | |
| additive GL | | | | -1.38 | -3.01 – 0.26 | 0.098 |
| **Random Effects** | | | | | | |
| $\sigma^2$ | 1.16 | | | 1.16 | | |
| $\tau_{00}$ | 0.18 $_{block}$ | | | 0.18 $_{block}$ | | |
| | 0.54 $_{gene\_pool}$ | | | 0.54 $_{gene\_pool}$ | | |
| ICC | 0.38 | | | 0.38 | | |
| N | 8 $_{block}$ | | | 8 $_{block}$ | | |
| | 6 $_{gene\_pool}$ | | | 6 $_{gene\_pool}$ | | |
| Observations | 1727 | | | 1727 | | |
| Marg / Cond $R^2$ | 0.000 / 0.381 | | | 0.001 / 0.382 | | |

| | $\delta^{13}C$ | | | $\delta^{13}C$ | | |
|---|---|---|---|---|---|---|
| *Predictors* | *Estimates* | *CI* | *p* | *Estimates* | *CI* | *p* |
| (Intercept) | -26.42 | -27.33 – -25.51 | **<0.001** | -25.87 | -27.05 – -24.69 | **<0.001** |
| recessive GL | -0.35 | -1.43 – 0.73 | 0.522 | | | |
| additive GL | | | | -1.41 | -3.26 – 0.44 | 0.136 |
| **Random Effects** | | | | | | |
| $\sigma^2$ | 1.05 | | | 1.05 | | |
| $\tau_{00}$ | 0.11 clone:prov | | | 0.11 clone:prov | | |
| | 0.01 altitude:gene_pool | | | 0.00 altitude:gene_pool | | |
| | 0.00 prov | | | 0.01 prov | | |
| | 0.18 block | | | 0.18 block | | |
| | 0.56 gene_pool | | | 0.56 gene_pool | | |
| ICC | 0.45 | | | 0.45 | | |
| N | 8 block | | | 8 block | | |
| | 458 clone | | | 458 clone | | |
| | 33 prov | | | 33 prov | | |
| | 33 altitude | | | 33 altitude | | |
| | 6 gene_pool | | | 6 gene_pool | | |
| Observations | 1723 | | | 1723 | | |
| Marg / Cond $R^2$ | 0.000 / 0.451 | | | 0.001 / 0.452 | | |

| | height_cm | | | height_cm | | |
|---|---|---|---|---|---|---|
| *Predictors* | *Estimates* | *CI* | *p* | *Estimates* | *CI* | *p* |
| (Intercept) | 378.91 | 170.58 – 587.24 | **<0.001** | 401.95 | 184.93 – 618.97 | **<0.001** |
| recessive GL | -94.31 | -178.85 – -9.77 | **0.029** | | | |
| additive GL | | | | -133.42 | -276.96 – 10.13 | 0.069 |
| **Random Effects** | | | | | | |
| $\sigma^2$ | 16609.24 | | | 16608.91 | | |
| $\tau_{00}$ | 1582.53 $_{clone:prov}$ | | | 1591.91 $_{clone:prov}$ | | |
| | 259.50 $_{block:site}$ | | | 259.76 $_{block:site}$ | | |
| | 739.53 $_{prov}$ | | | 734.73 $_{prov}$ | | |
| | 1683.93 $_{gene\_pool}$ | | | 1682.96 $_{gene\_pool}$ | | |
| | 52708.21 $_{site}$ | | | 52695.15 $_{site}$ | | |
| ICC | 0.77 | | | 0.77 | | |
| N | 40 $_{block}$ | | | 40 $_{block}$ | | |
| | 5 $_{site}$ | | | 5 $_{site}$ | | |
| | 464 $_{clone}$ | | | 464 $_{clone}$ | | |
| | 34 $_{prov}$ | | | 34 $_{prov}$ | | |
| | 6 $_{gene\_pool}$ | | | 6 $_{gene\_pool}$ | | |
| Observations | 11029 | | | 11029 | | |
| Marg / Cond $R^2$ | 0.000 / 0.774 | | | 0.000 / 0.774 | | |

| | height_cm | | | height_cm | | |
|---|---|---|---|---|---|---|
| *Predictors* | *Estimates* | *CI* | *p* | *Estimates* | *CI* | *p* |
| (Intercept) | 377.45 | 168.30 – 586.61 | **<0.001** | 400.83 | 183.08 – 618.59 | **<0.001** |
| recessive GL | -95.80 | -180.44 – -11.17 | **0.027** | | | |
| additive GL | | | | -135.25 | -278.93 – 8.44 | 0.065 |
| **Random Effects** | | | | | | |
| $\sigma^2$ | 16601.36 | | | 16601.02 | | |
| $\tau_{00}$ | 1584.10 $_{clone:prov}$ | | | 1593.70 $_{clone:prov}$ | | |
| | 259.60 $_{block:site}$ | | | 259.87 $_{block:site}$ | | |
| | 109.41 $_{altitude:gene\_pool}$ | | | 0.28 $_{altitude:gene\_pool}$ | | |
| | 609.04 $_{prov}$ | | | 715.85 $_{prov}$ | | |
| | 1999.81 $_{gene\_pool}$ | | | 1979.05 $_{gene\_pool}$ | | |
| | 52891.38 $_{site}$ | | | 52857.76 $_{site}$ | | |
| ICC | 0.78 | | | 0.78 | | |
| N | 40 $_{block}$ | | | 40 $_{block}$ | | |
| | 5 $_{site}$ | | | 5 $_{site}$ | | |
| | 463 $_{clone}$ | | | 463 $_{clone}$ | | |
| | 33 $_{prov}$ | | | 33 $_{prov}$ | | |
| | 33 $_{altitude}$ | | | 33 $_{altitude}$ | | |
| | 6 $_{gene\_pool}$ | | | 6 $_{gene\_pool}$ | | |
| Observations | 11003 | | | 11003 | | |
| Marg / Cond $R^2$ | 0.000 / 0.776 | | | 0.000 / 0.776 | | |

| | bud-burst | | | bud-burst | | |
|---|---|---|---|---|---|---|
| *Predictors* | *Estimates* | *CI* | *p* | *Estimates* | *CI* | *p* |
| (Intercept) | 1282.04 | 1236.48 – 1327.61 | **<0.001** | 1251.65 | 1176.85 – 1326.44 | **<0.001** |
| recessive GL | -7.62 | -88.16 – 72.91 | 0.853 | | | |
| additive GL | | | | 52.85 | -85.94 – 191.64 | 0.455 |
| **Random Effects** | | | | | | |
| $\sigma^2$ | 5894.04 | | | 5890.69 | | |
| $\tau_{00}$ | 791.62 $_{gene\_pool}$ | | | 828.54 $_{gene\_pool}$ | | |
| | 47.53 $_{block}$ | | | 46.88 $_{block}$ | | |
| ICC | 0.12 | | | 0.13 | | |
| N | 5 $_{block}$ | | | 5 $_{block}$ | | |
| | 6 $_{gene\_pool}$ | | | 6 $_{gene\_pool}$ | | |
| Observations | 1240 | | | 1240 | | |
| Marg / Cond $R^2$ | 0.000 / 0.125 | | | 0.000 / 0.130 | | |

| | bud-burst | | | bud-burst | | |
|---|---|---|---|---|---|---|
| *Predictors* | *Estimates* | *CI* | *p* | *Estimates* | *CI* | *p* |
| (Intercept) | 1287.61 | 1235.60 – 1339.62 | **<0.001** | 1263.41 | 1174.12 – 1352.71 | **<0.001** |
| recessive GL | -8.17 | -109.46 – 93.12 | 0.874 | | | |
| additive GL | | | | 40.08 | -130.97 – 211.12 | 0.646 |
| **Random Effects** | | | | | | |
| $\sigma^2$ | 4567.29 | | | 4568.43 | | |
| $\tau_{00}$ | 1143.97 $_{clone:prov}$ | | | 1138.43 $_{clone:prov}$ | | |
| | 230.56 $_{altitude:gene\_pool}$ | | | 232.19 $_{altitude:gene\_pool}$ | | |
| | 0.00 $_{prov}$ | | | 0.02 $_{prov}$ | | |
| | 485.84 $_{gene\_pool}$ | | | 513.39 $_{gene\_pool}$ | | |
| | 60.75 $_{block}$ | | | 60.36 $_{block}$ | | |
| ICC | 0.30 | | | 0.30 | | |
| N | 5 $_{block}$ | | | 5 $_{block}$ | | |
| | 360 $_{clone}$ | | | 360 $_{clone}$ | | |
| | 33 $_{prov}$ | | | 33 $_{prov}$ | | |
| | 33 $_{altitude}$ | | | 33 $_{altitude}$ | | |
| | 6 $_{gene\_pool}$ | | | 6 $_{gene\_pool}$ | | |
| Observations | 1238 | | | 1238 | | |
| Marg / Cond $R^2$ | 0.000 / 0.296 | | | 0.000 / 0.299 | | |

| | survival | | | survival | | |
|---|---|---|---|---|---|---|
| *Predictors* | *Odds Ratios* | *CI* | *p* | *Odds Ratios* | *CI* | *p* |
| (Intercept) | 4.64 | 0.42 – 50.80 | 0.209 | 5.21 | 0.39 – 68.97 | 0.211 |
| recessive GL | 0.51 | 0.15 – 1.77 | 0.287 | | | |
| additive GL | | | | 0.42 | 0.05 – 3.66 | 0.435 |
| **Random Effects** | | | | | | |
| $\sigma^2$ | 3.29 | | | 3.29 | | |
| $\tau_{00}$ | 0.18 $_{clone:prov}$ | | | 0.18 $_{clone:prov}$ | | |
| | 0.41 $_{block:site}$ | | | 0.41 $_{block:site}$ | | |
| | 0.09 $_{prov}$ | | | 0.09 $_{prov}$ | | |
| | 0.01 $_{gene\_pool}$ | | | 0.01 $_{gene\_pool}$ | | |
| | 7.00 $_{site}$ | | | 7.00 $_{site}$ | | |
| ICC | 0.70 | | | 0.70 | | |
| N | 40 $_{block}$ | | | 40 $_{block}$ | | |
| | 5 $_{site}$ | | | 5 $_{site}$ | | |
| | 464 $_{clone}$ | | | 464 $_{clone}$ | | |
| | 34 $_{prov}$ | | | 34 $_{prov}$ | | |
| | 6 $_{gene\_pool}$ | | | 6 $_{gene\_pool}$ | | |
| Observations | 17833 | | | 17833 | | |
| Marg / Cond $R^2$ | 0.000 / 0.700 | | | 0.000 / 0.700 | | |

| | survival | | | survival | | |
|---|---|---|---|---|---|---|
| *Predictors* | | *Odds Ratios* | | | *CI* | |
| (Intercept) | 4.66 | 0.46 – 47.64 | *p* | *Odds Ratios* | *CI* | *p* |
| recessive GL | 0.48 | 0.17 – 1.31 | 0.194 | 5.51 | 0.48 – 62.86 | 0.169 |
| additive GL | | | 0.150 | | | |
| | | | | 0.36 | 0.06 – 2.00 | 0.244 |
| **Random Effects** | | | | | | |
| $\sigma^2$ | 3.29 | | | 3.29 | | |
| $\tau_{00}$ | 0.40 $_{block:site}$ | | | 0.40 $_{block:site}$ | | |
| | 0.10 $_{altitude:gene\_pool}$ | | | 0.10 $_{altitude:gene\_pool}$ | | |
| | 0.01 $_{gene\_pool}$ | | | 0.01 $_{gene\_pool}$ | | |
| | 6.67 $_{site}$ | | | 6.67 $_{site}$ | | |
| ICC | 0.69 | | | 0.69 | | |
| N | 40 $_{block}$ | | | 40 $_{block}$ | | |
| | 5 $_{site}$ | | | 5 $_{site}$ | | |
| | 33 $_{altitude}$ | | | 33 $_{altitude}$ | | |
| | 6 $_{gene\_pool}$ | | | 6 $_{gene\_pool}$ | | |
| Observations | 17793 | | | 17793 | | |
| Marg / Cond $R^2$ | 0.000 / 0.685 | | | 0.000 / 0.685 | | |

**Table S3**. Linear mixed models of the deleteriousness scores based on Provean annotations with phenotypic traits related to water-use efficiency, tree growth and bud-burst in maritime pine for the two regional datasets from Corsica, including or not altitude of the provenance locality as random effect: (A) *CORSICA-capture* and (B) *CORSICA-array,* as well as for (C) the global dataset *GLOBAL-array*. For the latter dataset correlation with survival was also tested. The recessive Provean scores (recessive PR) load focus on average deleteriousness of homozygous sites containing deleterious mutations whereas the additive Provean scores (additive PR) indicates the average deleteriousness of deleterious alleles in homozygous and heterozygous sites. Traits used were: carbon isotope discrimination ($\delta^{13}C$) for water-use efficiency, tree height in cm for tree growth, and Julian day of entry in accumulated degree-days from the first day of the year for bud-burst. CI, confidence intervals; *p*, *p*-value based on conditional F-tests with Kenward-Roger approximation for the degrees of freedom; $\sigma^2$, residual variance; $\tau_{00}$, random effect variances; ICC, intraclass correlation coefficient; N, number of random effects groups; Marg. $R^2$, marginal goodness of fit (fixed effects); Cond. $R^2$, conditional goodness of fit (fixed and random effects).

A. *CORSICA-capture* dataset

| | | $\delta^{13}C$ | | | $\delta^{13}C$ | |
|---|---|---|---|---|---|---|
| *Predictors* | *Estimates* | *CI* | *p* | *Estimates* | *CI* | *p* |
| (Intercept) | -27.89 | -29.03 – -26.75 | **<0.001** | -27.02 | -29.01 – -25.03 | **<0.001** |
| Recessive PR | 11.15 | -1.41 – 23.71 | 0.081 | | | |
| Additive PR | | | | 12.36 | -0.92 – 25.65 | 0.068 |
| **Random Effects** | | | | | | |
| $\sigma^2$ | 0.45 | | | 0.45 | | |
| $\tau_{00}$ | 0.07 block:plot_id | | | 0.06 block:plot_id | | |
| | 0.06 plot_id | | | 0.06 plot_id | | |
| ICC | 0.23 | | | 0.22 | | |
| N | 3 block | | | 3 block | | |
| | 3 plot_id | | | 3 plot_id | | |
| Observations | 93 | | | 93 | | |
| Marg / Cond $R^2$ | 0.026 / 0.252 | | | 0.029 / 0.243 | | |

| | height_cm | | | height_cm | | |
|---|---|---|---|---|---|---|
| *Predictors* | *Estimates* | *CI* | *p* | *Estimates* | *CI* | *p* |
| (Intercept) | 262.77 | 211.94 – 313.61 | **<0.001** | 219.66 | 127.21 – 312.11 | **<0.001** |
| Recessive PR | 978.88 | 421.04 – 1536.72 | **0.001** | | | |
| Additive PR | | | | 268.94 | -350.46 – 888.33 | 0.393 |
| **Random Effects** | | | | | | |
| $\sigma^2$ | 1678.39 | | | 1793.70 | | |
| $\tau_{00}$ | 336.58 $_{block}$ | | | 340.72 $_{block}$ | | |
| ICC | 0.17 | | | 0.16 | | |
| N | 5 $_{block}$ | | | 5 $_{block}$ | | |
| Observations | 168 | | | 168 | | |
| Marg / Cond $R^2$ | 0.057 / 0.214 | | | 0.004 / 0.163 | | |

| | height_cm | | | height_cm | | |
|---|---|---|---|---|---|---|
| *Predictors* | *Estimates* | *CI* | *p* | *Estimates* | *CI* | *p* |
| (Intercept) | 258.26 | 207.50 – 309.02 | **<0.001** | 211.20 | 121.41 – 300.99 | **<0.001** |
| Recessive PR | 890.95 | 324.36 – 1457.54 | **0.002** | | | |
| Additive PR | | | | 193.11 | -410.44 – 796.67 | 0.528 |
| **Random Effects** | | | | | | |
| $\sigma^2$ | 1535.39 | | | 1615.55 | | |
| $\tau_{00}$ | 229.45 $_{altitude}$ | | | 266.20 $_{altitude}$ | | |
| | 158.54 $_{block}$ | | | 164.03 $_{block}$ | | |
| ICC | 0.20 | | | 0.21 | | |
| N | 5 $_{block}$ | | | 5 $_{block}$ | | |
| | 14 $_{altitude}$ | | | 14 $_{altitude}$ | | |
| Observations | 168 | | | 168 | | |
| Marg / Cond $R^2$ | 0.050 / 0.241 | | | 0.002 / 0.212 | | |

| | bud-burst | | | bud-burst | | |
|---|---|---|---|---|---|---|
| (Intercept) | 1990.95 | 1847.92 – 2133.98 | **<0.001** | 2065.14 | 1827.00 – 2303.29 | **<0.001** |
| Recessive PR | -161.69 | -1688.02 – 1364.65 | 0.834 | | | |
| Additive PR | | | | 413.58 | -1160.93 – 1988.10 | 0.604 |
| **Random Effects** | | | | | | |
| $\sigma^2$ | 10230.91 | | | 10209.52 | | |
| $\tau_{00}$ | 2298.07 $_{plot\_id}$ | | | 2371.28 $_{plot\_id}$ | | |
| ICC | 0.18 | | | 0.19 | | |
| N | 3 $_{plot\_id}$ | | | 3 $_{plot\_id}$ | | |
| Observations | 139 | | | 139 | | |
| Marg / Cond $R^2$ | 0.000 / 0.184 | | | 0.002 / 0.190 | | |

| | bud-burst | | | bud-burst | | |
|---|---|---|---|---|---|---|
| *Predictors* | *Estimates* | *CI* | *p* | *Estimates* | *CI* | *p* |
| (Intercept) | 1996.74 | 1846.66 – 2146.82 | **<0.001** | 2101.69 | 1863.02 – 2340.36 | **<0.001** |
| Recessive PR | -96.00 | -1681.25 – 1489.26 | 0.905 | | | |
| Additive PR | | | | 661.25 | -907.49 – 2229.98 | 0.406 |
| **Random Effects** | | | | | | |
| $\sigma^2$ | 9668.47 | | | 9569.61 | | |
| $\tau_{00}$ | 732.27 $_{altitude}$ | | | 832.97 $_{altitude}$ | | |
| | 2380.18 $_{plot\_id}$ | | | 2552.41 $_{plot\_id}$ | | |
| ICC | 0.24 | | | 0.26 | | |
| N | 3 $_{plot\_id}$ | | | 3 $_{plot\_id}$ | | |
| | 11 $_{altitude}$ | | | 11 $_{altitude}$ | | |
| Observations | 139 | | | 139 | | |
| Marg / Cond $R^2$ | 0.000 / 0.244 | | | 0.004 / 0.264 | | |

B. *CORSICA-array* dataset

| | δ13C | | | δ13C | | |
|---|---|---|---|---|---|---|
| *Predictors* | *Estimates* | *CI* | *p* | *Estimates* | *CI* | *p* |
| (Intercept) | -29.01 | -30.01 – -28.00 | **<0.001** | -29.09 | -30.06 – -28.11 | **<0.001** |
| Recessive PR | -0.65 | -2.55 – 1.24 | 0.497 | | | |
| Additive PR | | | | -0.99 | -4.04 – 2.05 | 0.519 |
| **Random Effects** | | | | | | |
| $\sigma^2$ | 0.34 | | | 0.34 | | |
| $\tau_{00}$ | 0.25 plot_id | | | 0.26 plot_id | | |
| ICC | 0.42 | | | 0.43 | | |
| N | 3 plot_id | | | 3 plot_id | | |
| Observations | 120 | | | 120 | | |
| Marg. / Cond. $R^2$ | 0.002 / 0.426 | | | 0.002 / 0.429 | | |

| | height_cm | | | height_cm | | |
|---|---|---|---|---|---|---|
| *Predictors* | *Estimates* | *CI* | *p* | *Estimates* | *CI* | *p* |
| (Intercept) | 205.82 | 185.62 – 226.02 | **<0.001** | 207.20 | 177.68 – 236.71 | **<0.001** |
| Recessive PR | 79.51 | -4.98 – 164.01 | 0.065 | | | |
| Additive PR | | | | 78.04 | -57.94 – 214.03 | 0.260 |
| **Random Effects** | | | | | | |
| $\sigma^2$ | 1901.97 | | | 1913.81 | | |
| $\tau_{00}$ | 128.04 block:plot_id | | | 132.65 block:plot_id | | |
| | 58.37 plot_id | | | 57.94 plot_id | | |
| ICC | 0.09 | | | 0.09 | | |
| N | 3 block | | | 3 block | | |
| | 3 plot_id | | | 3 plot_id | | |
| Observations | 334 | | | 334 | | |
| Marg / Cond $R^2$ | 0.009 / 0.098 | | | 0.004 / 0.094 | | |

| | height_cm | | | height_cm | | |
|---|---|---|---|---|---|---|
| Predictors | Estimates | CI | p | Estimates | CI | p |
| (Intercept) | 205.13 | 184.36 – 225.90 | **<0.001** | 209.18 | 178.96 – 239.39 | **<0.001** |
| Recessive PR | 83.30 | -1.17 – 167.77 | 0.053 | | | |
| Additive PR | | | | 95.23 | -41.29 – 231.76 | 0.171 |
| **Random Effects** | | | | | | |
| $\sigma^2$ | 1797.75 | | | 1806.22 | | |
| $\tau_{00}$ | 126.34 $_{altitude}$ | | | 129.43 $_{altitude}$ | | |
| | 63.06 $_{block:plot\_id}$ | | | 66.76 $_{block:plot\_id}$ | | |
| | 98.58 $_{plot\_id}$ | | | 103.06 $_{plot\_id}$ | | |
| ICC | 0.14 | | | 0.14 | | |
| N | 3 $_{block}$ | | | 3 $_{block}$ | | |
| | 3 $_{plot\_id}$ | | | 3 $_{plot\_id}$ | | |
| | 22 $_{altitude}$ | | | 22 $_{altitude}$ | | |
| Observations | 334 | | | 334 | | |
| Marg / Cond $R^2$ | 0.026 / 0.143 | | | 0.024 / 0.143 | | |

| | bud_burst | | | bud_burst | | |
|---|---|---|---|---|---|---|
| *Predictors* | *Estimates* | *CI* | *p* | *Estimates* | *CI* | *p* |
| (Intercept) | 1980.85 | 1907.51 – 2054.18 | **<0.001** | 2050.46 | 1948.58 – 2152.34 | **<0.001** |
| Recessive PR | -130.51 | -431.57 – 170.55 | 0.393 | | | |
| Additive PR | | | | 249.33 | -237.44 – 736.09 | 0.313 |
| **Random Effects** | | | | | | |
| $\sigma^2$ | 10804.43 | | | 10792.83 | | |
| $\tau_{00}$ | 97.08 $_{block:plot\_id}$ | | | 136.48 $_{block:plot\_id}$ | | |
| | 997.82 $_{plot\_id}$ | | | 824.13 $_{plot\_id}$ | | |
| ICC | 0.09 | | | 0.08 | | |
| N | 3 $_{block}$ | | | 3 $_{block}$ | | |
| | 3 $_{plot\_id}$ | | | 3 $_{plot\_id}$ | | |
| Observations | 153 | | | 153 | | |
| Marg / Cond $R^2$ | 0.004 / 0.096 | | | 0.006 / 0.088 | | |

| | bud_burst | | | bud_burst | | |
|---|---|---|---|---|---|---|
| *Predictors* | *Estimates* | *CI* | *p* | *Estimates* | *CI* | *p* |
| Intercept) | 1979.34 | 1903.71 – 2054.97 | **<0.001** | 2047.62 | 1941.40 – 2153.84 | **<0.001** |
| Recessive PR | -147.27 | -458.10 – 163.55 | 0.351 | | | |
| Additive PR | | | | 230.69 | -282.49 – 743.87 | 0.375 |
| **Random Effects** | | | | | | |
| $\sigma^2$ | 10629.35 | | | 10730.11 | | |
| $\tau_{00}$ | 291.61 $_{altitude}$ | | | 108.67 $_{altitude}$ | | |
| | 24.86 $_{block:plot\_id}$ | | | 98.85 $_{block:plot\_id}$ | | |
| | 1007.00 $_{plot\_id}$ | | | 831.69 $_{plot\_id}$ | | |
| ICC | 0.11 | | | 0.09 | | |
| N | 3 $_{block}$ | | | 3 $_{block}$ | | |
| | 3 $_{plot\_id}$ | | | 3 $_{plot\_id}$ | | |
| Observations | 153 | | | 153 | | |
| Marg / Cond $R^2$ | 0.006 / 0.116 | | | 0.006 / 0.093 | | |

C. *GLOBAL-array* dataset

| | $\delta^{13}C$ | | | $\delta^{13}C$ | | |
|---|---|---|---|---|---|---|
| (Intercept) | -26.47 | -27.13 – -25.81 | **<0.001** | -26.47 | -27.20 – -25.75 | **<0.001** |
| Recessive PR | 0.61 | -0.38 – 1.60 | 0.226 | | | |
| Additive PR | | | | 0.52 | -1.24 – 2.28 | 0.563 |
| **Random Effects** | | | | | | |
| $\sigma^2$ | 1.05 | | | 1.05 | | |
| $\tau_{00}$ | 0.11 clon:prov | | | 0.11 clon:prov | | |
| | 0.01 prov | | | 0.01 prov | | |
| | 0.18 block | | | 0.18 block | | |
| | 0.48 gene_pool_k8 | | | 0.50 gene_pool_k8 | | |
| ICC | 0.43 | | | 0.43 | | |
| N | 8 block | | | 8 block | | |
| | 459 clon | | | 459 clon | | |
| | 34 prov | | | 34 prov | | |
| | 6 gene_pool_k8 | | | 6 gene_pool_k8 | | |
| $\sigma^2$ | 1.05 | | | 1.05 | | |
| $\tau_{00}$ | 0.11 clon:prov | | | 0.11 clon:prov | | |
| Observations | 1723 | | | 1723 | | |
| Marg / Cond $R^2$ | 0.001 / 0.429 | | | 0.000 / 0.433 | | |

| | $\delta^{13}C$ | | | $\delta^{13}C$ | | |
|---|---|---|---|---|---|---|
| *Predictors* | *Estimates* | *CI* | *p* | *Estimates* | *CI* | *p* |
| (Intercept) | -26.49 | -27.17 – -25.81 | **<0.001** | -26.49 | -27.24 – -25.75 | **<0.001** |
| Recessive PR | 0.58 | -0.42 – 1.57 | 0.254 | | | |
| Additive PR | | | | 0.48 | -1.28 – 2.24 | 0.593 |
| **Random Effects** | | | | | | |
| $\sigma^2$ | 1.05 | | | 1.05 | | |
| $\tau_{00}$ | 0.11 $_{clon:prov}$ | | | 0.11 $_{clon:prov}$ | | |
| | 0.01 $_{altitude:gene\_pool\_k8}$ | | | 0.01 $_{altitude:gene\_pool\_k8}$ | | |
| | 0.00 $_{prov}$ | | | 0.00 $_{prov}$ | | |
| | 0.18 $_{block}$ | | | 0.18 $_{block}$ | | |
| | 0.53 $_{gene\_pool\_k8}$ | | | 0.54 $_{gene\_pool\_k8}$ | | |
| ICC | 0.44 | | | 0.45 | | |
| N | 8 $_{block}$ | | | 8 $_{block}$ | | |
| | 458 $_{clon}$ | | | 458 $_{clon}$ | | |
| | 33 $_{prov}$ | | | 33 $_{prov}$ | | |
| | 33 $_{altitude}$ | | | 33 $_{altitude}$ | | |
| | 6 $_{gene\_pool\_k8}$ | | | 6 $_{gene\_pool\_k8}$ | | |
| Observations | 1723 | | | 1723 | | |
| Marg / Cond $R^2$ | 0.001 / 0.442 | | | 0.000 / 0.447 | | |

| | height_cm | | | height_cm | | |
|---|---|---|---|---|---|---|
| *Predictors* | *Estimates* | *CI* | *p* | *Estimates* | *CI* | *p* |
| (Intercept) | 333.14 | 128.30 – 537.99 | **0.001** | 300.88 | 94.78–506.98 | **0.004** |
| Recessive PR | -6.16 | -77.36–65.04 | 0.865 | | | |
| Additive PR | | | | -178.61 | -303.68 –53.54 | **0.005** |
| **Random Effects** | | | | | | |
| $\sigma^2$ | 16609.66 | | | 16603.54 | | |
| $\tau_{00}$ | 1607.74 $_{clon:prov}$ | | | 1587.19 $_{clon:prov}$ | | |
| | 259.80 $_{block:site}$ | | | 260.26 $_{block:site}$ | | |
| | 724.27 $_{prov}$ | | | 712.20 $_{prov}$ | | |
| | 1732.65 $_{gene\_pool\_k8}$ | | | 1884.21 $_{gene\_pool\_k8}$ | | |
| | 52727.30 $_{site}$ | | | 52734.92 $_{site}$ | | |
| ICC | 0.77 | | | 0.77 | | |
| N | 40 $_{block}$ | | | 40 $_{block}$ | | |
| | 5 $_{site}$ | | | 5 $_{site}$ | | |
| | 464 $_{clon}$ | | | 464 $_{clon}$ | | |
| | 34 $_{prov}$ | | | 34 $_{prov}$ | | |
| | 6 $_{gene\_pool\_k8}$ | | | 6 $_{gene\_pool\_k8}$ | | |
| Observations | 11029 | | | 11029 | | |
| Marg / Cond $R^2$ | 0.000 / 0.775 | | | 0.000 / 0.774 | | |

| | height_cm | | | height_cm | | |
|---|---|---|---|---|---|---|
| *Predictors* | *Estimates* | *CI* | *p* | *Estimates* | *CI* | *p* |
| (Intercept) | 329.03 | 123.50 – 534.57 | **0.002** | 304.36 | 97.27 – 511.45 | **0.004** |
| Recessive PR | -19.84 | -98.19 – 58.51 | 0.620 | | | |
| Additive PR | | | | -149.90 | -288.34 – -11.47 | **0.034** |
| **Random Effects** | | | | | | |
| $\sigma^2$ | 16601.31 | | | 16601.71 | | |
| $\tau_{00}$ | 1610.41 clon:prov | | | 1587.30 clon:prov | | |
| | 259.95 block:site | | | 260.05 block:site | | |
| | 0.26 altitude:gene_pool_k8 | | | 611.24 altitude:gene_pool_k8 | | |
| | 705.86 prov | | | 80.54 prov | | |
| | 2032.90 gene_pool_k8 | | | 2179.95 gene_pool_k8 | | |
| | 52784.84 site | | | 52856.77 site | | |
| ICC | 0.78 | | | 0.78 | | |
| N | 40 block | | | 40 block | | |
| | 5 site | | | 5 site | | |
| | 463 clon | | | 463 clon | | |
| | 33 prov | | | 33 prov | | |
| | 33 altitude | | | 33 altitude | | |
| | 6 gene_pool_k8 | | | 6 gene_pool_k8 | | |
| Observations | 11003 | | | 11003 | | |
| Marg / Cond $R^2$ | 0.000 / 0.776 | | | 0.000 / 0.776 | | |

| | bud-burst | | | bud-burst | | |
|---|---|---|---|---|---|---|
| *Predictors* | *Estimates* | *CI* | *p* | *Estimates* | *CI* | *p* |
| (Intercept) | 1287.93 | 1262.17 – 1313.69 | **<0.001** | 1301.78 | 1265.20 – 1338.37 | **<0.001** |
| Recessive PR | 24.76 | -68.65 – 118.17 | 0.603 | | | |
| Additive PR | | | | 98.30 | -65.01 – 261.61 | 0.238 |
| **Random Effects** | | | | | | |
| $\sigma^2$ | 4561.06 | | | 4562.17 | | |
| $\tau_{00}$ | 1144.74 $_{clon:prov}$ | | | 1144.41 $_{clon:prov}$ | | |
| | 219.03 $_{prov}$ | | | 195.94 $_{prov}$ | | |
| | 498.53 $_{gene\_pool\_k8}$ | | | 506.42 $_{gene\_pool\_k8}$ | | |
| | 60.46 $_{block}$ | | | 61.19 $_{block}$ | | |
| ICC | 0.30 | | | 0.29 | | |
| N | 5 $_{block}$ | | | 5 $_{block}$ | | |
| | 361 $_{clon}$ | | | 361 $_{clon}$ | | |
| | 34 $_{prov}$ | | | 34 $_{prov}$ | | |
| | 6 $_{gene\_pool\_k8}$ | | | 6 $_{gene\_pool\_k8}$ | | |
| Observations | 1240 | | | 1240 | | |
| Marg / Cond $R^2$ | 0.000 / 0.297 | | | 0.002 / 0.296 | | |

| | bud-burst | | | bud-burst | | |
|---|---|---|---|---|---|---|
| *Predictors* | *Estimates* | *CI* | *p* | *Estimates* | *CI* | *p* |
| (Intercept) | 1287.43 | 1261.61 – 1313.25 | **<0.001** | 1301.41 | 1264.82 – 1338.01 | **<0.001** |
| Recessive PR | 24.34 | -69.14 – 117.83 | 0.610 | | | |
| Additive PR | | | | 98.71 | -64.74 – 262.16 | 0.236 |
| **Random Effects** | | | | | | |
| $\sigma^2$ | 4567.09 | | | 4568.08 | | |
| $\tau_{00}$ | 1146.00 $_{\text{clon:prov}}$ | | | 1145.83 $_{\text{clon:prov}}$ | | |
| | 215.74 $_{\text{altitude:gene\_pool\_k8}}$ | | | 197.79 $_{\text{altitude:gene\_pool\_k8}}$ | | |
| | 5.38 $_{\text{prov}}$ | | | 0.00 $_{\text{prov}}$ | | |
| | 496.61 $_{\text{gene\_pool\_k8}}$ | | | 502.60 $_{\text{gene\_pool\_k8}}$ | | |
| | 60.85 $_{\text{block}}$ | | | 61.63 $_{\text{block}}$ | | |
| ICC | 0.30 | | | 0.29 | | |
| N | 5 $_{\text{block}}$ | | | 5 $_{\text{block}}$ | | |
| | 360 $_{\text{clon}}$ | | | 360 $_{\text{clon}}$ | | |
| | 33 $_{\text{prov}}$ | | | 33 $_{\text{prov}}$ | | |
| | 33 $_{\text{altitude}}$ | | | 33 $_{\text{altitude}}$ | | |
| | 6 $_{\text{gene\_pool\_k8}}$ | | | 6 $_{\text{gene\_pool\_k8}}$ | | |
| Observations | 1238 | | | 1238 | | |
| Marg / Cond $R^2$ | 0.000 / 0.297 | | | 0.002 / 0.296 | | |

| | survival | | | survival | | |
|---|---|---|---|---|---|---|
| *Predictors* | *Odds Ratios* | *CI* | *p* | *Odds Ratios* | *CI* | *p* |
| (Intercept) | 3.33 | 0.32 – 34.30 | 0.313 | 3.19 | 0.30 – 33.76 | 0.335 |
| Recessive PR | 0.91 | 0.29 – 2.88 | 0.876 | | | |
| Additive PR | | | | 0.73 | 0.10 – 5.54 | 0.762 |
| **Random Effects** | | | | | | |
| $\sigma^2$ | 3.29 | | | 3.29 | | |
| $\tau_{00}$ | 0.18 $_{clon:prov}$ | | | 0.18 $_{clon:prov}$ | | |
| | 0.41 $_{block:site}$ | | | 0.41 $_{block:site}$ | | |
| | 0.09 $_{prov}$ | | | 0.09 $_{prov}$ | | |
| | 0.00 $_{gene\_pool\_k8}$ | | | 0.00 $_{gene\_pool\_k8}$ | | |
| | 6.99 $_{site}$ | | | 6.99 $_{site}$ | | |
| ICC | 0.70 | | | 0.70 | | |
| N | 40 $_{block}$ | | | 40 $_{block}$ | | |
| | 5 $_{site}$ | | | 5 $_{site}$ | | |
| | 464 $_{clon}$ | | | 464 $_{clon}$ | | |
| | 34 $_{prov}$ | | | 34 $_{prov}$ | | |
| | 6 $_{gene\_pool\_k8}$ | | | 6 $_{gene\_pool\_k8}$ | | |
| Observations | 17833 | | | 17833 | | |
| Marg / Cond $R^2$ | 0.000 / 0.700 | | | 0.000 / 0.700 | | |

| | survival | | | survival | | |
|---|---|---|---|---|---|---|
| *Predictors* | *Odds Ratios* | *CI* | *p* | *Odds Ratios* | *CI* | *p* |
| (Intercept) | 3.32 | 0.32 – 34.01 | 0.312 | 3.19 | 0.30 – 33.46 | 0.334 |
| Recessive PR | 0.91 | 0.29 – 2.87 | 0.871 | | | |
| Additive PR | | | | 0.73 | 0.10 – 5.59 | 0.766 |
| **Random Effects** | | | | | | |
| $\sigma^2$ | 3.29 | | | 3.29 | | |
| $\tau_{00}$ | 0.18 $_{clon:prov}$ | | | 0.18 $_{clon:prov}$ | | |
| | 0.41 $_{block:site}$ | | | 0.41 $_{block:site}$ | | |
| | 0.09 $_{altitude:gene\_pool\_k8}$ | | | 0.09 $_{altitude:gene\_pool\_k8}$ | | |
| | 0.00 $_{prov}$ | | | 0.00 $_{prov}$ | | |
| | 0.00 $_{gene\_pool\_k8}$ | | | 0.00 $_{gene\_pool\_k8}$ | | |
| | 6.99 $_{site}$ | | | 6.99 $_{site}$ | | |
| ICC | 0.70 | | | 0.70 | | |
| N | 40 $_{block}$ | | | 40 $_{block}$ | | |
| | 5 $_{site}$ | | | 5 $_{site}$ | | |
| Observations | 17793 | | | 17793 | | |
| Marg / Cond $R^2$ | 0.000 / 0.700 | | | 0.000 / 0.700 | | |
